# Hierarchical cross-scale analysis identifies parallel ventral striatal networks coding for dynamic and stabilized olfactory reward predictions

**DOI:** 10.1101/2021.02.22.432268

**Authors:** Laurens Winkelmeier, Carla Filosa, Max Scheller, Renée Hartig, Markus Sack, Robert Becker, David Wolf, Jonathan Reinwald, Martin Fungisai Gerchen, Alexander Sartorius, Andreas Meyer-Lindenberg, Wolfgang Weber-Fahr, Christian Clemm von Hohenberg, Eleonora Russo, Wolfgang Kelsch

## Abstract

The unbiased identification of brain circuits responsible for behavior and their local cellular computations is a challenge for neuroscience. We establish here a hierarchical cross-scale approach from behavioral modeling and fMRI in task-performing mice to cellular network dynamics to identify how reward predictions are represented in the forebrain upon olfactory conditioning. fMRI identified functional segregation in reward prediction and error computations among olfactory cortices and subcortical circuits. Among them, the olfactory tubercle contributed both to dynamic reward predictions and prediction error. In this region, cellular recordings revealed two parallel neuronal populations for prediction coding. One population produced stabilized predictions as distributed stimulus-bound transient network activity; the other evolved during anticipatory waiting and fully reflected predicted value in single-units, dynamically integrating the recent cue-specific history of uncertain outcomes. Thus, the cross-scale approach revealed regional functional differentiation among the distributed forebrain circuits with a limbic hotspot for multiple non-redundant reward prediction coding.

## INTRODUCTION

Complex functions like stimulus-outcome learning involve a chain of cognitive processes comprising the recognition of unexpected outcomes and the updating of reward predictions (RP) (Dayan and Niv, 2008, Schultz, 2000). These processes are thought to be computed across distributed human forebrain networks (Daw et al., 2005, De Martino et al., 2006, Ferenczi et al., 2016, Li et al., 2011, O’Doherty et al., 2004). While enormous insights have emerged concerning a limited number of intensely studied brain regions in small rodents (Cohen et al., 2012, Sul et al., 2010, Takahashi et al., 2016, Tian et al., 2016), it remains incompletely clear whether rodents employ similar distributed circuits for reward prediction error computations that have been identified by whole-brain imaging in humans.

A more unbiased systems perspective in translational models would require structured approaches that parametrize behavior and identify brain regions functionally involved. The identified circuits can then be examined at finer granularity to disentangle the underlying neural codes and circuit mechanisms. Within such a hierarchical approach, fMRI is an attractive tool to assess, at mesoscopic scale and in an unsupervised fashion, the contributions of cortical and subcortical regions to task computations. Recent developments demonstrate the potential of fMRI for dissecting reward and fear circuits in awake, passive mice (Harris et al., 2015, Lee et al., 2016) or connectivity during resting state in mouse mutants (Tsurugizawa et al., 2020). Even though fMRI has become a standard tool to assess task-related brain activity in humans, to date, only very few fMRI studies exist in rodents during task performance (Han et al., 2019).

Here we developed a hierarchical approach, from mesoscale fMRI to cellular recordings, to dissect olfactory reward prediction coding in the context of stimulus-outcome learning. During such reinforcement learning, an initially neutral sensory stimulus (called a conditioned stimulus; CS) is paired repeatedly after a waiting interval with reward (an unconditioned stimulus; US) (Flagel et al., 2011, Oettl et al., 2020, Schultz, 2000, Setlow et al., 2003, Takahashi et al., 2016). Midbrain dopamine neurons (DAN) show firing responses to the CS and US. In crude terms, these responses code a temporal difference prediction error (PE) signal, which computes the mismatch between the value of the received reward and the RP (Montague et al., 1996, Schultz et al., 1997). These signals change with learning. Initially, when stimulus-outcome associations are uncertain, DAN mainly fire bursts at US, reflecting the positive surprise about the obtained rewards (Schultz et al., 1997). Then, during learning, an additional response evolves at CS, reflecting the reward probability associated with the sensory stimulus (Fiorillo et al., 2003). A pioneering study in mice examined how the RP and PE are formed in DAN by their monosynaptic inputs from subcortical regions (Tian et al., 2016). Nonetheless, much of the connectivity in the distributed cortical and subcortical reward networks is polysynaptic (Ikemoto et al., 2015), and an unbiased assessment of the distributed RP computations in the rodent forebrain is still missing. Specifically, it is not entirely clear to which extent the different olfactory cortices, the attached higher regions, as well as the subcortical circuits, contribute respectively to the multiple facets of value estimation.

To assess the differential contributions of olfactory forebrain regions to the distributed RP and PE computations, mice were trained in an olfactory stimulus-outcome learning task with different reward probabilities. We first modeled the behavior to formulate specific predictions about RP and PE signals during the task and regress them on fMRI data from mice performing the task in the scanner to identify key forebrain regions. We then studied the underlying computations with single-unit recordings in two fMRI-identified cortical and striatal regions. With this hierarchical approach, we revealed functional differentiation of olfactory cortices in value coding and multiple non-redundant parallel computations of the RP in an fMRI-identified region, the olfactory tubercle of the ventral striatum.

## RESULTS

### Reward prediction and its updating by the recent outcome-history

To disentangle the regions involved in distributed computations of reward prediction and error in the mouse forebrain, a cohort of 23 animals, examined later with fMRI, was habituated to a head-fixed set-up and trained on stimulus-outcome pair associations. Mice were conditioned to odor stimuli that signaled different probabilities of upcoming reward (100%, 50%, or 0%, hereafter these trial types are labeled CS100, CS50, or CS0, respectively) (**Fig. 1A**). The odor stimulus was presented for 1 s followed by a 1.7 s wait interval before the water drop was delivered with the given reward probability (**Fig. 1A**). The water port was positioned so that mice had to actively lick to sense and retrieve the reward. After training, mice performed above criterion and licked in more than 80% of CS100 and CS50 trials, but not in the CS0 trials (**Fig. S1A**). Lick intensities in the waiting window correlated with the probability of reward to that odor cue (**Fig. 1B-C**). In 10 trained mice, we monitored pupil responses as a second proxy for the animals’ reward expectation (**Fig. 1D**; see also **Fig S1B,C**). Pupil dilation equally reflected the stimulus-specific RP (**Fig. 1E**). Thus, mice learned to predict reward outcomes upon CS presentation.

**Figure 1.**
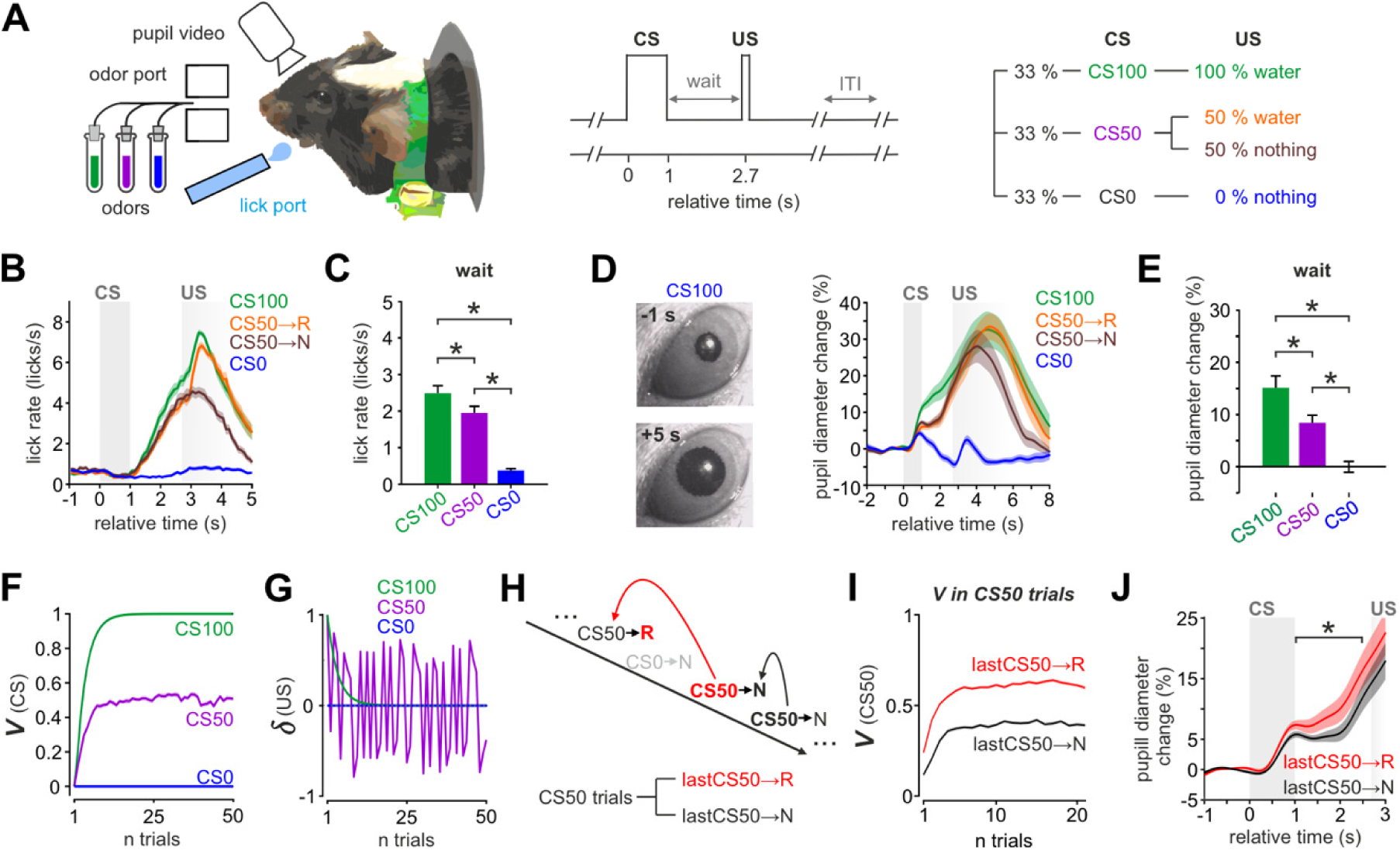
Trained mice display differential anticipatory responses to olfactory stimulus-outcome pairs and dynamic value update by recent reward history. (**A**) Mice were trained to learn stimulus-outcome pairs: Three different odors (CS) were applied for 1 s followed by a fixed waiting period of 1.7 s before a drop of water was delivered (US). Licking behavior and pupil diameter were monitored simultaneously. Stimulus CS100 and CS50 predicted US with different reward probabilities (100% and 50%). Stimulus CS0 was never rewarded. (**B**) Evolution of average lick rates ± SEM differentiated trial types (n = 69 sessions in 23 animals). (**C**) Average lick rate ± SEM in the waiting window differentiated the respective trial types with CS100>CS50>CS0 (one-way ANOVA with Tukey post-hoc comparisons). (**D**) Video frames illustrating pupil image 1 s before and 5 s after onset of CS100 (left). Average change ± SEM of the baseline subtracted pupil diameter also differentiated trial types (right; n = 10 sessions in 10 animals). (**E**) The average change ± SEM of the baseline subtracted pupil diameter revealed similar pupil responses in the waiting period to the lick responses (one-way ANOVA with Tukey post-hoc comparisons). (**F**) Averaged RP values *V*(*CS*) of the three CS from the TD model (n = 100 simulated CS-US sequences, learning rate *α* = 0.28 optimized from pupillary data, initial conditions set to zero). The values approached the true reward probabilities associated with each CS. (**G**) The TD PE *δ* at US approached zero in CS100 and CS0 trials, while it remained large in CS50 trials as shown for an exemplary session. **(H)** Scheme illustrating the subdivision of CS50 trials for recent outcome-history analysis. CS50 trials were grouped according to the outcome of the previous CS50 trial (rewarded: lastCS50→R; unrewarded: lastCS50→N). **(I)** Average modeled *V*(*CS*) for CS50 trials divided according to lastCS50→R and lastCS50→N predicts a dynamic update of RP based on the recent outcome-history. **(J)** Average change ± SEM of the normalized pupil diameter in CS50 trials split for lastCS50→R and lastCS50→N. The change in pupil diameter during waiting followed the recent history update (two-tailed paired t-test). In the figure: * indicates p < 0.05 (see Table S1 for exact p-values and test details) and n indicates the number of units.

We used reinforcement learning modeling to parametrize the RP value associated at CS with each odor (‘*V*(*CS*)’), and the PE at US (‘*δ*(*US*)’). We built a temporal difference (TD) model with the learning rate (‘*α*’) set as a free parameter and optimized on the pupil responses of the behavioral sessions. The average of the learning rates was used to build a TD model applied throughout the study. Exploiting the generative power of the TD model, we simulated 100 sessions and observed that the average of *V*(*CS*) across sessions approached the true reward frequencies associated with each CS (**Fig. 1F**). As expected, the absolute value of *δ* at the time of potential reward became small in CS100 and CS0, but remained large for CS50 trials (**Fig. 1G**). In CS50 trials, the reward probability was at chance level throughout the session. Yet, the RP value was dynamically updated at each trial to integrate local fluctuations on the outcome probabilities. According to the TD model, the RP associated with a CS at each time instance *t* results from the integration of the whole history of rewards paired (or not) with each instance of CS. Each obtained reward will increase the RP associated with CS50 while the absence of reward will diminish it. Consistently, when dividing CS50 trials according to the outcome of the preceding CS50 trial, the recent outcome-history was reflected in the state values of the TD model (**Fig. 1H-I**) and confirmed behaviorally by the pupil responses during the waiting interval (**Fig. 1J**). Thus, animals learned to predict reward outcome at CS modulating RP by the recent outcome-history of uncertain rewards as predicted by the TD model.

### Distributed forebrain representations of the reward prediction error

We employed fMRI to localize forebrain regions involved in the olfactory RP, its modulation by the recent outcome-history, and the PE. A Bruker 9.4T rodent MR-scanner was used to image mice during task performance. A mouse MRI cradle was designed with odor and lick ports, connectors for head bars, and a cover over the back of the mouse (**Fig. 2A**). Mice were habituated to the cradle and head fixation in mock scanners, progressively adding the task paradigm, and then increasing levels of MRI sound recorded during an fMRI session. After completion of this training (**Fig. S1D**), the cohort of 23 mice underwent fMRI. Each individual performed up to four sessions in the scanner on separate days. To reduce stress and assure task performance, mice performed 20-30 preparatory trials before commencing the BOLD imaging sequence. In order to maintain comparable levels of satiety and motivation as the 150-trial-sessions outside the scanner, the task-related session comprised only 120 trials following the preparatory trials. Sessions where animals did not lick at reward were stopped and the animals were not used for further sessions. Of the 67 completed scanning sessions, 51 sessions in 18 animals performed above criterion (> 80% correct hits and rejections per session) (**Fig. S1E-G**).

**Figure 2.**
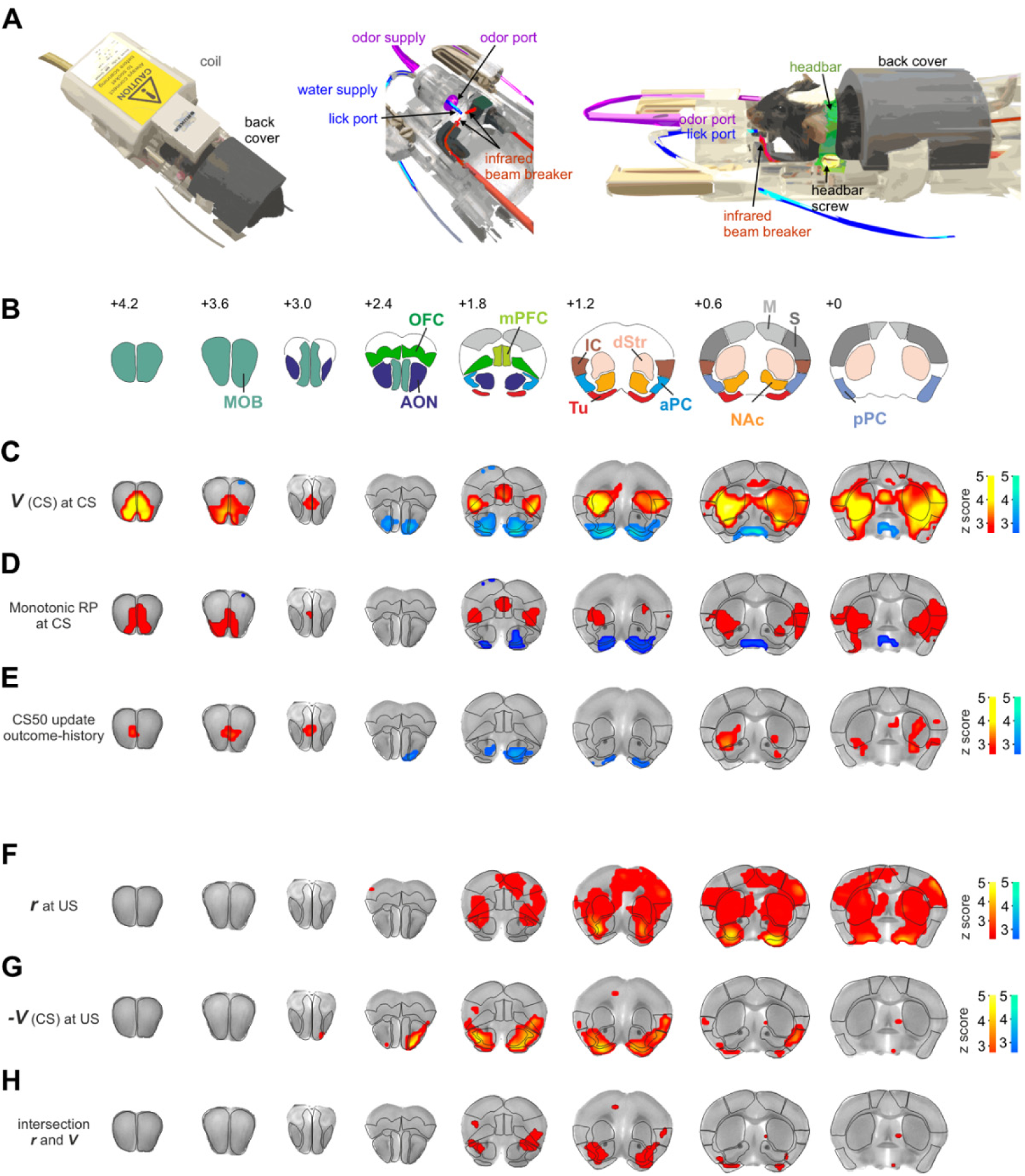
Distributed brain circuits code for monotonic reward prediction and the prediction error. (**A**) Illustration of the MRI compatible behavioral setup with MRI coil (left) comprises an odor port, a lick port, respective supply tubes, an infrared beam breaker, a head-fixation system, and a back cover (middle and right). (**B**) Anatomical regions of interest (ROI) definitions based on Paxinos atlas (location from Bregma indicated in mm). Abbreviations: anterior olfactory nucleus (AON), anterior piriform cortex (aPC), dorsal striatum (dStr), insular cortex (IC), main olfactory bulb (MOB), medial prefrontal cortex (mPFC), motor cortex (M), nucleus accumbens (NAc), olfactory tubercle (Tu), orbitofrontal cortex (OFC), posterior piriform cortex (pPC), sensory cortex (S). (**C**) Group-level Z-statistical maps showing BOLD correlates of RP values *V*(*CS*) from the TD model (n = 51 sessions in 18 animals). Statistical threshold was set to p < 0.025 false discovery rate (FDR)-corrected, for two-sided testing (as for the other maps unless otherwise indicated). Red color spectrum indicates areas with positive correlations to RP values, while blue colors indicate negative correlations. The regression revealed recruitment of olfactory and striatal brain regions with opposing effects on BOLD. Note that amongst the sections shown, the postero-ventral parts of pPC have signal dropout preventing detection of (de-) activation. (**D**) Within the regions associated with *V*(*CS*) from (C), monotonic RP (CS100>CS50>CS0, red or CS100<CS50<CS0, blue) was represented mainly in prefrontal, insular, striatal regions, and MOB. Maps were created by intersecting contrast maps CS100>CS50 and CS50>CS0, each thresholded at p < 0.025 FDR-corrected; and vice versa: CS100<CS50 and CS50<CS0 (see the respective contrast maps in Fig. S2H-I). (**E**) Recent outcome-history modulating the BOLD response to CS50. Within the *V*(*CS*) regions from (C), we tested for a CS50 response enhancement by the reward in the previous CS50 trial. In olfactory and striatal regions, the CS50-associated response was indeed strengthened by a positive outcome-history. **(F-H)** In the TD model, the PE at US is *r* – *V*(*CS*). We identified brain regions correlating with PE by intersecting the regressor maps of (F) *r* and (G) *-V(CS)* at US, each thresholded at p < 0.05 FDR-corrected. The intersection, shown in (H), was largely confined to the lateral NAc, the posterior Tu, the AON, and lateral OFC.

We imaged a larger olfactory network comprising the main olfactory bulb (MOB), the primary olfactory and prefrontal cortices, and the ventral and dorsal striatum (**Fig. 2B**). Upon preprocessing (see Methods section and **Fig. S2A-C**), we computed a general linear model (GLM) on the BOLD time series of the 51 sessions, modeling CS and US timepoints as events, which were parametrically modulated as described below, and convolved with a previously determined mouse hemodynamic response function (Lebhardt et al., 2016). When parametrically modulating the CS regressor with the model-estimated RP value at CS, *V*(*CS*), we observed broad recruitment of brain regions (**Fig. 2C**) with positive signals in the MOB, the prefrontal cortices, specifically the medial prefrontal cortex (mPFC) and the orbitofrontal cortex (OFC), the insular cortex, and parts of the posterior piriform cortex (pPC). Negative BOLD signals to *V*(*CS*) were found in the olfactory primary cortices, namely the anterior olfactory nucleus (AON) and partially also the anterior piriform cortex (aPC). The dorsal striatum and the two ventral striatal brain regions, consisting of the olfactory tubercle (Tu) and nucleus accumbens (NAc), were also correlated to *V*(*CS*).

Importantly, the applied GLM is not primarily designed to distinguish between a monotonic computation (CS100 > CS50 > C0) of *V*(*CS*) and a simple binary differentiation between presence and absence of reward expectation (e.g., CS100 = CS50 > CS0). Indeed, CS100 and CS50 recruited similar brain regions, unlike CS0 (**Fig. S2D-G**). Therefore, we computed a separate GLM where the three CS types were modelled with individual regressors, and we determined monotonic RP correlates by the intersection of CS contrasts (CS100 > CS50 and CS50 > CS0, or vice versa for negative contrasts, **Fig. S2H,I**). Within the *V*(*CS*)-significant regions, monotonic RP was expressed primarily in a network comprising prefrontal regions (mPFC, OFC, anterior insular cortex), striatal circuits (Tu, parts of NAc and lateral striatum) as well as primary olfactory regions, namely the MOB-AON loop and partially the pPC (**Fig. 2D**). This monotonic RP network was largely determined by the narrow contrast between the two rewarded stimuli CS100 and CS50 (**Fig. S2H**). Again, the AON and Tu displayed monotonic RP with negative contrasts. We then investigated, among the regions representing *V*(*CS*), which brain regions participated in the RP updating based on the recent outcome-history, as predicted by the TD model (cf. Fig. 1H-J). To this end, the CS50 regressor was parametrically modulated by the binary outcome of the preceding CS50 trial. Among the regions representing *V*(*CS*), the three striatal regions and the MOB-AON loop reflected the cue-specific outcome-history updating (**Fig. 2E**).

In the TD model, the PE *δ* at US is *r* - *V*(*CS*). In humans, forebrain regions contribute to either one of the two PE-at-US components (*r* or –*V*(*CS*)), and only few regions compute both (Hare et al., 2008, Niv et al., 2012). To test for BOLD correlates of the PE at US, we parametrically modulated the US regressor with –*V*(*CS*) and with *r* in the same GLM. We found correlates of *r* (**Fig. 2F**) in the lateral NAc, the posterior Tu, the dorsal striatum and, additionally, also in some cortices including the OFC, the insula, and, among the olfactory cortices, prominently the AON. In contrast, –*V*(*CS*) at US (**Fig. 2G**) involved broadly a ventral stream of olfactory and striatal regions and additionally the OFC. The intersection of *r* and –*V*(*CS*) (**Fig. 2H**) was prominent in the lateral NAc, but comprised also the posterior Tu, the AON and lateral OFC; thus, providing a relatively confined network involved in the PE at US for olfactory stimulus-outcome learning.

In summary, monotonic RP and the PE components were encoded in partially overlapping networks of olfactory, prefrontal and striatal brain circuits. Among them, few regions including the Tu also displayed RP updating by the recent cue-specific outcome-history. Notably, the contribution of the aPC to these computations was relatively restricted compared to other olfactory regions and mainly contributed to the PE value component.

### Task-related neuronal population coding in the olfactory cortex and striatum

fMRI had revealed unexpected contributions of the two olfactory brain circuits, aPC and Tu, to the value computations in the forebrain circuit. To dissect the local cellular coding underlying these distributed value computations, we performed single-unit’ recordings in a separate cohort of 11 mice with a chronic custom-designed tetrode array. Dual site recordings with up to 64 channels per brain region were performed in the aPC and Tu (**Fig. 3A**; **Fig. S3A-E**). We examined trained animals that performed the task above criterion (**Fig. S3F-H**) and obtained 169 putative striatal projection neurons with baseline firing below 5 Hz in the Tu (**Fig. S3E**). Additionally, we obtained 486 regular-firing neurons in the aPC with baseline firing below 10 Hz. To capture coherent positive and negative changes in the population firing rate, we computed the deflection from baseline of the population vector during the trial (**Fig. 3B**). We computed the Euclidean distance between the population vector at each time instance and at baseline, to detect changes in angle or rate of the population activity. Tu population activity coded for the predicted reward value of the stimulus at CS and during the waiting period (**Fig. 3C-D**). We quantified then to which extent different odors and outcomes recruited the same units. By computing the Pearson cross-correlation between the average instantaneous population vectors associated with different CS, we found progressive recruitment of units encoding cross-stimulus reward anticipation during the waiting period of CS50 and CS100 trials, and progressive decorrelation between the rewarded and non-rewarded CS trials (**Fig. 3E**). At US, population trajectories during CS50 trials diverged according to the outcome and, in case of reward, were pushed closer to the trajectory of CS100 trials (**Fig. 3F**).

**Figure 3.**
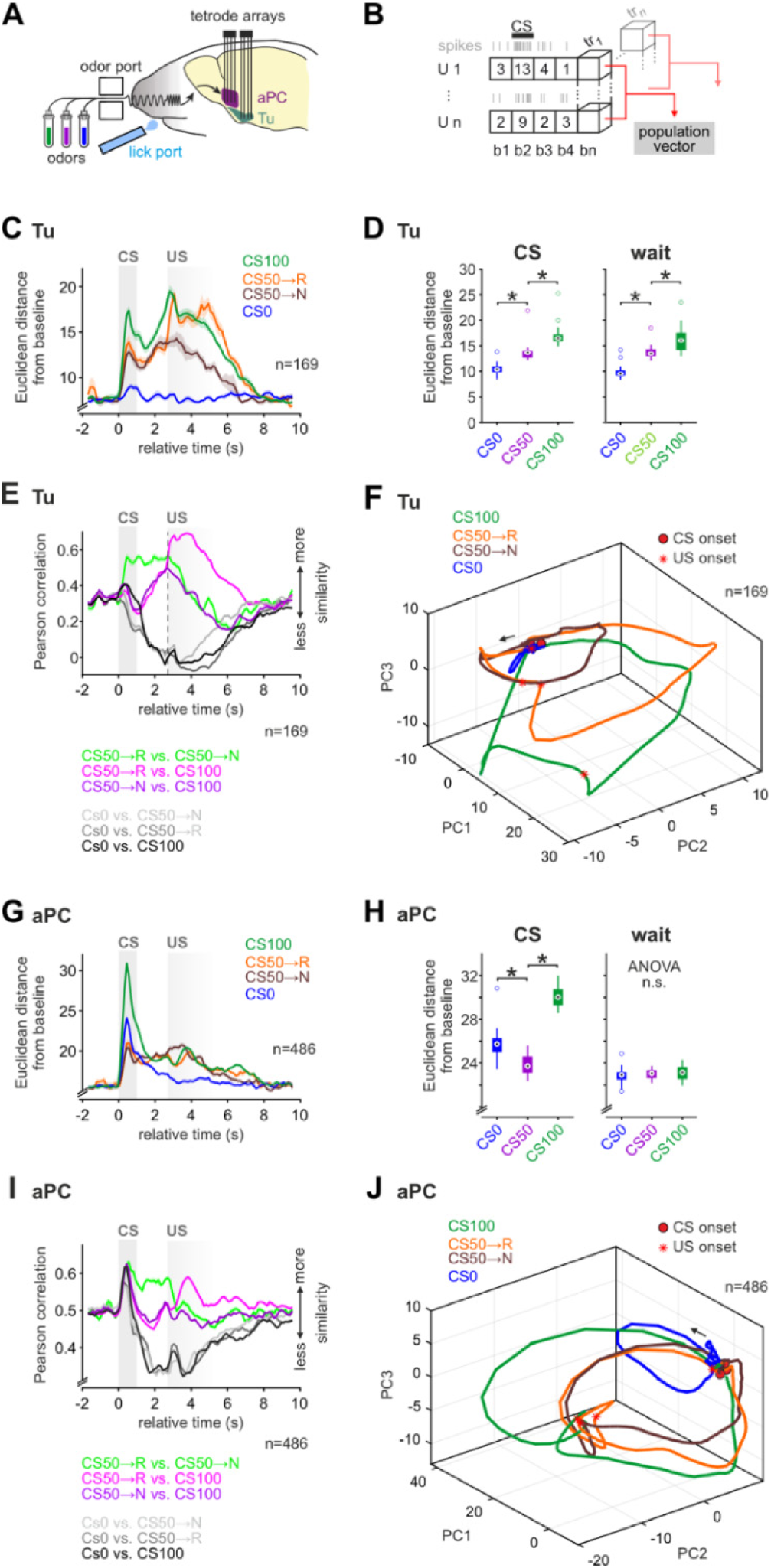
Task-related neuronal population coding in the olfactory cortex and striatum. (**A**) Configurations of the dual-site single-unit recordings. Custom-made tetrode arrays were implanted into the Tu and aPC. Mice performed the same stimulus-outcome associated task as in head-fixed configuration outside the scanner. (**B**) Population vectors were constructed by concatenating the firing rate of individual single units during bins of 250 ms and pooled across sessions. (**C**) The Euclidean distance of Tu population vectors from baseline displayed an RP coding at CS and during the waiting period, as well as reward coding at US. Displayed mean ± SEM, 500 ms bins slid in steps of 125 ms. (**D**) Euclidean distance from baseline during CS (from 0 to 1 s) and waiting window (from 1 to 2.5 s). One-way ANOVA with Tukey post hoc comparisons performed separately in each window. (**E**) Pearson correlation between Tu population vectors displayed as mean across pairs of trials ± SEM. The population responses to CS100 and CS50 became more similar during wait, suggesting a partial overlap in Tu units coding for the two stimuli. (**F**) First three principal components of the time-embedded averaged trajectories of Tu during different trial types. The arrow marks the direction of the temporal evolution. Reward delivery abruptly deflected the population trajectories of CS50→R and CS50→N. **(G-H)** Same as (C-D) for aPC. aPC population displayed a dominant response to all CS, but no monotonic RP coding. (**G**) Same as (E) for aPC. The similarity between aPC population response to all stimulus onsets is consistent with a sensory detection signal. (**H**) Same as (F) for aPC. Also aPC population trajectories reflect an initial indiscriminate sensory detection signal. In the figure: * indicates p < 0.05 (see Table S1 for exact p-values and test details) and n indicates the number of units.

The aPC displayed different population responses (**Fig. 3G-J**). In contrast to the Tu, aPC population activity was most pronounced during CS (**Fig. 3G**), as expected for a primary sensory cortex, and did not reflect RP (**Fig. 3G-H**). The CS onset triggered a correlated detection response across all trial types in aPC (**Fig. 3I**). This was also seen in the initial upshot of the population trajectories common to all stimuli (**Fig. 3J**, contrary to the Tu cf. Fig. 3F). Thus, while aPC had a pronounced detection response to olfactory stimuli, the Tu population coded for monotonic RP both during odor presentation as well as during the anticipatory waiting in trained animals.

### Task-inhibited value responses dominate aPC

We then aimed to better understand what cellular activity patterns underlie the negative BOLD correlates of value in the aPC. In trained mice, after an initial positive response to CS detection, negative mean population rate changes prevailed during waiting and upon US for all trial types (**Fig. 4A-B**). Interestingly, this negative rate change encoded predicted stimulus value during waiting, but not at CS, and responded to different reward outcomes at US (**Fig. 4B, right**). The forebrain imaging and the so far examined population coding were obtained from animals trained for at least two weeks in the specific task. As learning may modify the proportions between excited and inhibited responses (Bamford et al., 2018), we wondered whether the predominance of task-inhibited responses emerged with training. We thus analyzed the first session the animal ever faced the task with the three stimuli CS100, CS50 and CS0 (**Fig. 4C, S4A-C**). In this initial training session, the net population rate response was relatively balanced (**Fig. 4C**). With training, the fraction of units with task-excited responses did not change. Yet, their relative contribution in the CS50 and CS100 trials markedly dropped due to the increase of task-inhibited responses, especially at US (**Fig. S4D-E**).

**Figure 4.**
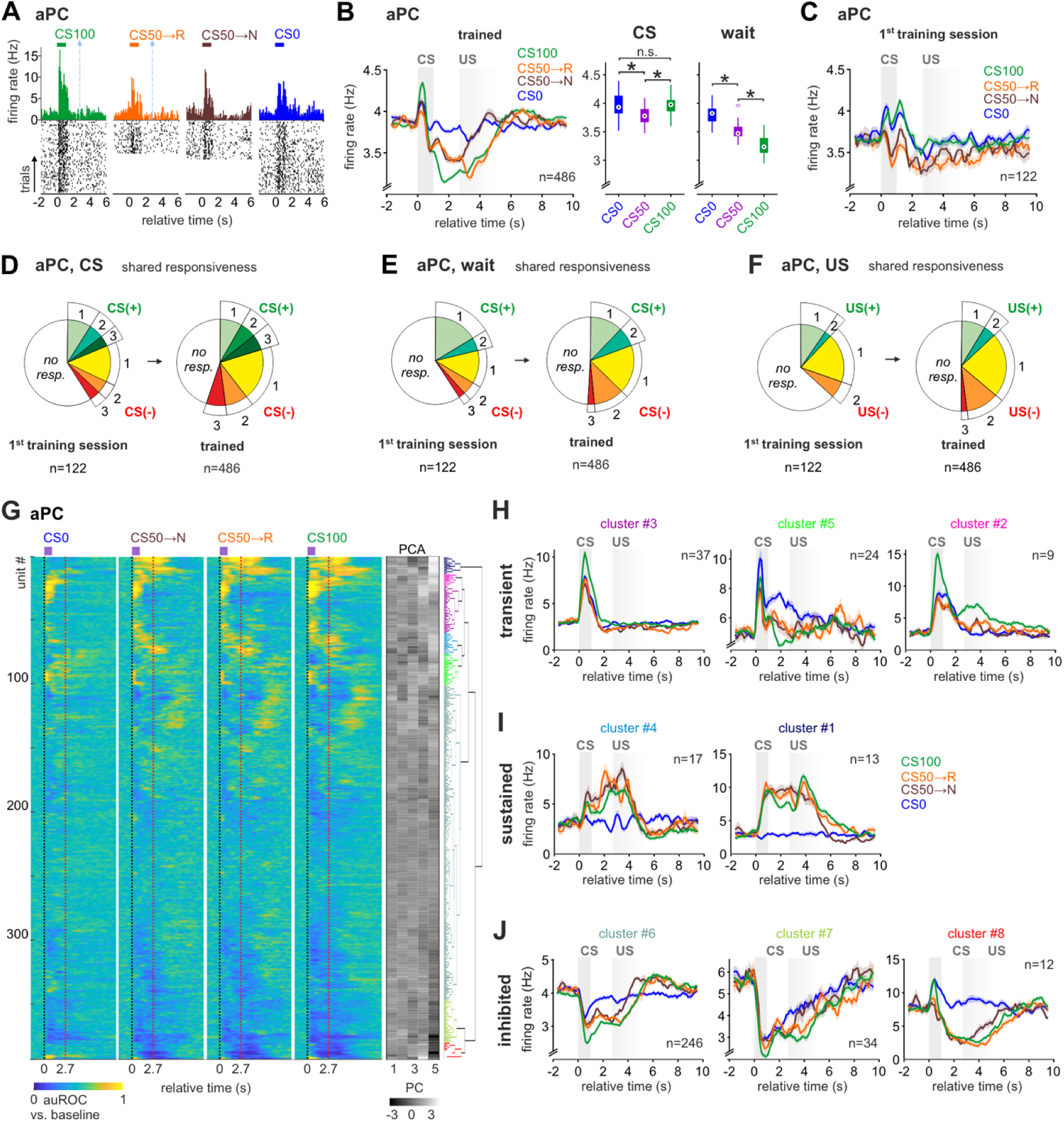
Task-inhibited value responses dominate aPC. (**A**) Example of single-unit firing activity in aPC in trained animals. Peri-stimulus time histogram (PSTH) and spike raster plots showed stable task-related responses. (**B**) Task-related mean firing rate evolution ± SEM in aPC units. After a first detection peak at CS, aPC displayed monotonic RP during the waiting period as a progressive decrease in firing rate (one-way ANOVA with Tukey post-hoc comparisons). (**C**) Box plots showing median aPC firing rate for CS and waiting period (one-way ANOVA with Tukey post-hoc comparisons). Solid bars indicate the 25^th^ and 75^th^ percentiles. **(D-F)** The fraction of either task-excited (+) or inhibited (-) responses to either one, two, or three trial types at (D) CS, (E) during wait, and (F) at US in the aPC. Unit responsiveness was tested against baseline with a Friedman test, p < 0.05 with Benjamini-Hochberg correction. The fraction of task-inhibited units increased after training together with their shared responsiveness to different CS. The fraction of excited units, as well as their level of selectivity to one or more trial types, remained relatively stable. **(G)** Classification of aPC units into functional clusters. Unit responsiveness during different trial types was quantified by the area under the receiver operating characteristic curve (auROC, left). The auROC was computed for each time bin by comparing the distribution of firing rates of the bin with that of baseline. The values 0, 0.5 and 1 indicate respectively lower, equal, and higher firing rate than during baseline, respectively. Hierarchical clustering (right) was performed on the first five principal components (center, gray-scale) of the auROC traces and revealed 8 prominent task-response clusters. **(H-J)** The task-response clusters identified in (G) were grouped according to their responsiveness at CS and during the waiting period. We identified three major groups with (H) task-excited transient activity at CS, (I) task-excited sustained activity during waiting, and (J) task-inhibited responses. Displayed mean firing rate ± SEM. The color of the titles matches the cluster color in the dendrogram in (G). Unlike the task-inhibited cluster, none of the task-excited clusters displayed a monotonic RP (see Fig. S4F-H). In the figure: * indicates p < 0.05 (see Table S1 for exact p-values and test details) and n indicates the number of units.

The development of value coding at the single-unit level would require an acquired generalization in the unit responses to rewarded stimuli. In line with a lack of value coding, aPC units with task-excited responses did not change their selectivity with learning (**Fig. 4D-F**). Compatible with inhibitory value coding, however, the number of units with task-inhibited responses to two stimuli increased in trained animals (**Fig. 4D-F**). Learning thus appeared to affect units negatively modulated by the task. To confirm this observation and account for a potential heterogeneity of aPC subpopulations contributing to RP coding, we clustered neurons of trained animals according to their task activation profiles (**Fig. 4G**). Units clustered in a variety of subpopulations and fell into three main groups according to their dominant characteristics. The main subpopulations consisted of transient CS-excited units (**Fig. 4H**) and a large group of task-inhibited units (**Fig. 4J**). Moreover, an additional smaller group of units displayed heterogenous sustained activity during waiting (**Fig. 4I**). Neither of the two task-excited cluster groups showed monotonic RP in their average response (**Fig. S4F-G**), which was, however, again reflected in the task-inhibited population (**Fig. 4J, S4H**). Thus, with training, aPC shifted towards task-inhibited responses that contributed to value coding.

### Parallel populations encode reward prediction in Tu

We then examined the cellular dynamics underlying the RP and PE correlates in the olfactory striatum revealed by fMRI (**Fig. 5**). In trained mice, the mean population firing responses encoded RP at CS and during waiting, as well as reward surprise at US (**Fig. 5A-B**). As expected, during the first session, reward surprise, but not RP, dominated task responses (**Fig. 5C**). Value coding began to emerge during the first session with the stimulus responses to the fully-rewarded CS100 differentiating from the other stimuli (**Fig. S5A**). Notably, the mean rate changes to CS0 were still positive in this first training session. With subsequent learning, the fraction of units with task-excited responses dropped in CS0 trials and increased in the rewarded trial types. As in aPC, training increased the fraction of units with task-inhibited responses across trial types (**Fig. S5B-C**). Contrary to aPC, the number of Tu units responding to more than one CS increased both for task-excited and task-inhibited responses (**Fig. 5D-F**). As expected for learned categorization of stimuli according to their predicted outcome value, the increase was caused by shared responsiveness to both CS50 and CS100 (**Fig. 5G-I**). This trend characterized all task epochs. However, we found some notable differences in the responsiveness of Tu units during the CS and the subsequent waiting interval. At CS, the total number of units with shared responsiveness to CS50 and CS100 was smaller than the sum of units responsive selectively to CS50 or CS100 (**Fig. 5G**). During waiting, instead, the large majority of recruited units shared responsiveness to CS50 and CS100 (**Fig. 5H**). This difference could suggest the presence of distinct functional Tu subpopulations active at different task epochs, and heterogeneity in the RP coding scheme during stimulus presentation and the subsequent anticipation of reward while waiting.

**Figure 5.**
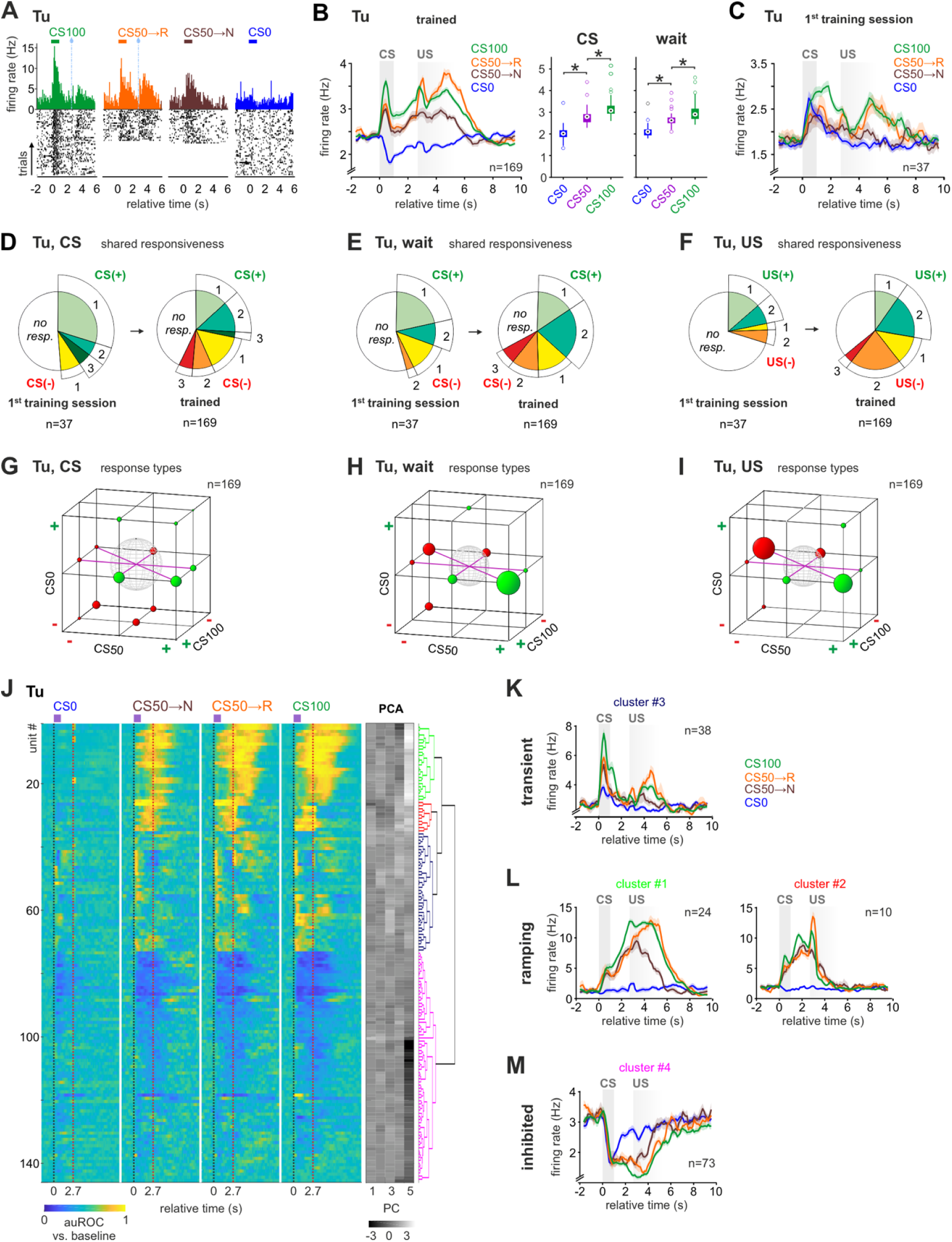
Tu units form transient and ramping task response clusters. **(A-C)** Same as Fig. 4A-C for Tu. After training, the task-related mean firing rate evolution ± SEM in Tu units reflected monotonic RP and reward surprise. **(D-F)** Same as Fig. 4D-F for Tu. The fraction of either task excited (+) or inhibited (-) responses in multiple SOP increased with training at (D) CS, (E) during wait, and (F) at US. **(G-I)** Fractions of selective and shared unit responses to different trial types (G) at CS, (H) during wait, and (I) at US in the Tu. The size of the spheres corresponds to the respective fraction of units. Green color indicates task-excited responses (+), while red color stands for task inhibited responses (-). Note the shared coding of rewarded trial types during wait and at US. **(J)** Same as Fig. 4G for Tu. In the Tu, PCA based hierarchical clustering revealed 4 task-response clusters. **(K-M)** Same as Fig. 4H-J for Tu. Mean firing rate ± SEM for each cluster identified in (J). According to their responsiveness at CS and during wait, single units were grouped into three functional clusters composed of units with: (K) task-excited, transient activity at CS, (L) task-excited, ramping activity during waiting, and (M) task-inhibited responses. Both the transient- and ramping-clusters displayed a monotonic RP (see Fig. S5D-F). In the figure: * indicates p < 0.05 (see Table S1 for exact p-values and test details) and n indicates the number of units.

To investigate the existence of distinct functional subpopulations in Tu, we clustered neurons of trained animals according to their task activation profiles (**Fig. 5J**). Tu units divided into three major groups. Task-excited units with transient activity at CS (transient-cluster, **Fig. 5K**), task-excited units with ramping activity during the waiting window (ramping-cluster, **Fig. 5L**), and units with task-inhibited responses (inhibited-cluster, **Fig. 5M**). The transient-cluster encoded monotonic RP at CS, which decayed shortly after the odor presentation had ceased (**Fig. 5K; Fig. S5D**). Within the ramping-clusters, instead, monotonic RP coding evolved only during waiting (**Fig. 5L; Fig. S5E**). In contrast to the task-excited clusters, the average firing rate of the inhibited-cluster did not significantly encode a monotonic RP (**Fig. 5M; Fig. S5F**). These findings suggest that stimulus-triggered and anticipatory RP were encoded by parallel task-excited neuronal populations in the Tu.

To better understand the coding mechanisms employed by such functionally distinct neuronal clusters, we examined whether the RP was computed in single neurons, whose rate response reflects the full set of reward probabilities, or only at the population level (**Fig. 6**). While more than half of the units in the ramping-clusters displayed a monotonic RP coding (**Fig. 6A-B**), in the transient-cluster only a quarter did so, and exclusively during odor presentation (**Fig. 6A-B, Fig. S6A**). Considering that the CS-bound value coding units are reinforced from pre-existing odor-specific responses (Oettl et al., 2020), we wondered whether the single-units not encoding monotonic RP still contributed collectively to it. Indeed, upon removal of all units with monotonic RP coding in the transient-cluster, the cumulative firing rate of the remaining units still reflected a robust population coding of the monotonic RP (**Fig. 6C**). This was not the case in the ramping-cluster (**Fig. S6B**). Thus, while the ramping-cluster encoded the full information at the single-unit level, the transient-cluster employed largely a distributed coding scheme. Only few task-inhibited units individually encoded a monotonic RP (**Fig. 6A**; see also **Fig. S6A**). In contrast to Tu, only a small fraction of all aPC units encoded monotonic RP above chance level (**Fig. S6C**), with the exception of a few units from the task-inhibited cluster. In fact, the occurrence of units with response intensities monotonically ordered (CS100 > CS50 > CS0 and CS0 > CS50 > CS100) passed chance level for all cluster groups in Tu (while no other intensity permutation did), but only for the task-inhibited cluster groups in aPC.

**Figure 6.**
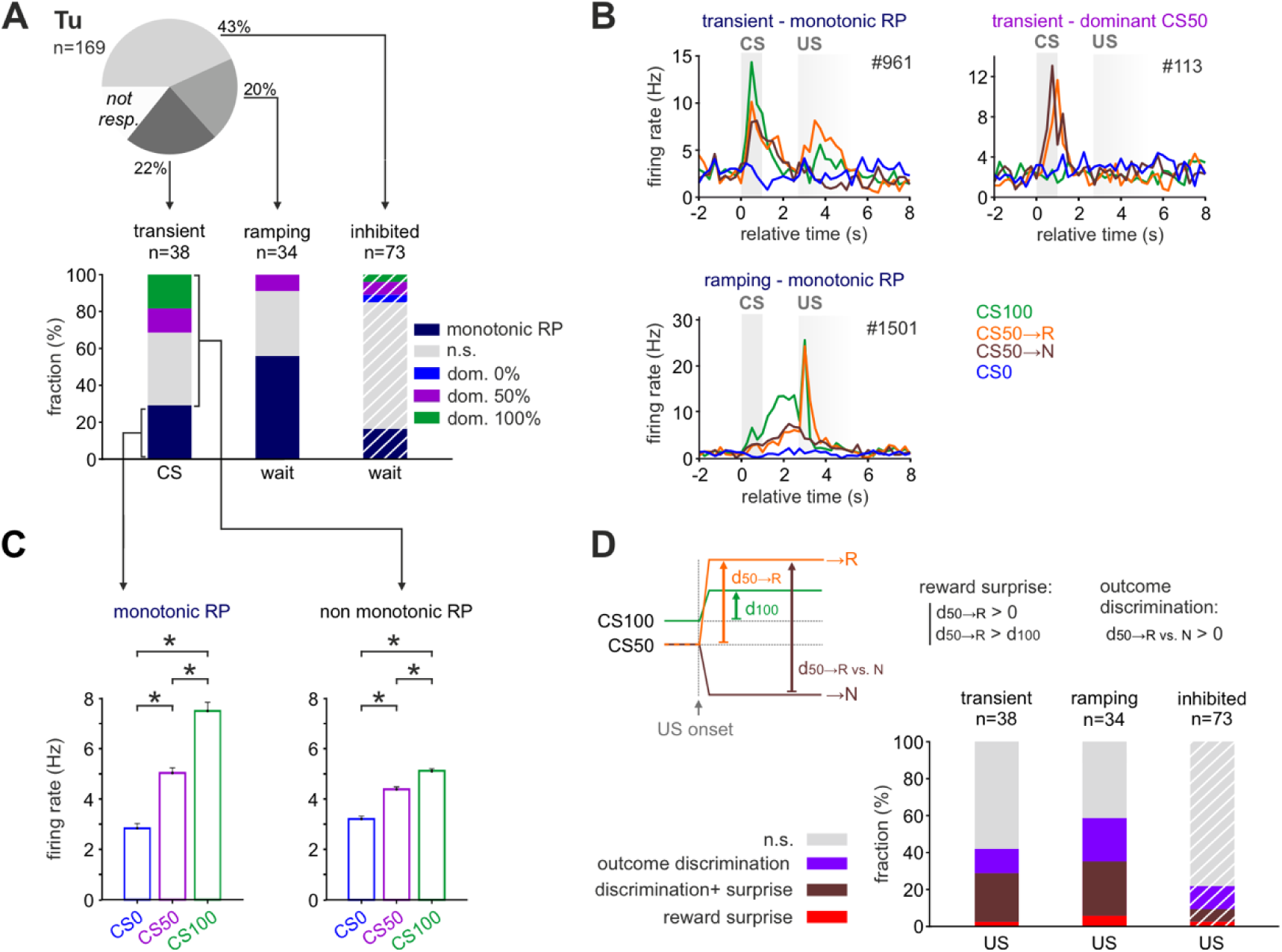
Transient and ramping units differently encode monotonic RP. (**A**) Pie chart (top) showing the percentage of the Tu units in the three major response clusters from Fig. 5K-M. The bar graphs (bottom) indicate the percentage of units in each cluster coding monotonic RP or showing a dominant response to one of the CS (the test window is indicated below each bar). Note that more than half of the units in the ramping-cluster displayed monotonic RP during wait while only a quarter from the transient-cluster did so at CS. Tests for monotonic RP and CS coding were performed via two-tailed t-test with significance threshold at p < 0.05. Dashed white lines indicate that the task-inhibited cluster was tested for reduction in firing rate (see Methods and Fig. S6A). (**B**) Average firing rate of example single units illustrating (upper left) monotonic RP coding or (upper right) dominant CS50 activation at CS in two units from the transient-cluster, and (lower left) monotonic RP coding during waiting of a unit in the ramping-cluster. (**C**) Average firing rate ± SEM at CS of the units in the transient-cluster that encoded monotonic RP individually (left) or not (right) (one-way ANOVA with Tukey post-hoc comparisons). Also units not individually coding monotonic RP displayed a robust monotonic RP as a population. Thus, the transient cluster displayed monotonic RP by distributed coding. For ramping-clusters see Fig. S6B. (**D**) Scheme illustrating reward surprise and outcome discrimination (top). Units were categorized as coding for reward surprise if during CS50 rewarded trials their increase in firing rate from before to after reward delivery (d_50→R_) was positive and bigger than that in CS100 trials (d_100_). Units were categorized as coding for outcome discrimination if the firing rate at US was higher for rewarded than for unrewarded trials (d_50→R vs. N_ > 0). Tests were performed via two-tailed t-test with significance threshold at p < 0.05. Dashed white lines indicate that the task-inhibited cluster was tested for reduction in firing rate (see Methods). (Bottom) Fraction of units from the three Tu clusters with reward surprise or outcome discrimination or both. In the figure: * indicates p < 0.05 (see Table S1 for exact p-values and test details) and n indicates the number of units.

To understand whether the Tu units with transient CS-bound value coding and those with ramping anticipation are differentially involved in PE computations at US, we tested if single-units encoded reward surprise and if they differentiated outcomes between reward and non-reward in CS50 trials (**Fig. 6D**). Reward surprise was defined by a larger rate change in rewarded CS50 than in CS100 trials from the waiting period to the US response. Note that negative surprise for unrewarded CS50 trials was not tested because phasic negative responses to US from the wait plateau cannot be distinguished from the progressive passive return to baseline. Approximately half of the units in both the transient- and ramping-clusters significantly encoded either reward surprise or outcome discrimination, with a large fraction of them significant for both discrimination and surprise. It is important to note that ramping neurons had a more sustained and pronounced response to reward than transient neurons (**Fig. 5K-L**), both relative to baseline (**Fig. S5D-E**) and to the anticipatory ramp (**Fig. S5G**). In the task-inhibited cluster, only a minority of units displayed surprise or outcome discrimination at US (**Fig. 6D**). In the aPC, roughly a quarter of units in all response clusters uniformely encoded outcome discrimination or reward surprise (**Fig. S6D**).

In summary, two parallel populations in the Tu - both encoding the PE - distinguished for the coding of RP in different task components: The CS-bound RP was computed with information distributed in single neurons responsive to specific CS, while anticipatory RP was computed as full information already in single neurons with ramping activity in anticipation of reward.

### Reward prediction is updated by its recent outcome-history

We finally examined how the multiple circuits encoding RP in the limbic system updated according to the recent outcome-history (**Fig. 7**). We divided the CS50 trials based on the outcome of the preceding CS50 trial (**Fig. 7A**). To compare this effect with the unselective effect of recent rewards associated with other CS (termed here ‘satiety’), we also examined the modulation of expected value in CS50 trials depending on whether the previous trial was CS100 or CS0 (**Fig. 7B**). We found that during waiting, anticipatory licking following CS50 was positively modulated if the previous CS was rewarded, integrating information of at least four prior CS50 trials (**Fig. 7C-E**), even though other trial types were interspersed (**Fig. S7A**). Satiety had, instead, the opposite effect on the licking behavior. The prior experience of either a CS0 or a CS100 trial increased or decreased, respectively, the intensity of anticipatory licking in CS50 trials (**Fig. 7C,F**).

**Figure 7.**
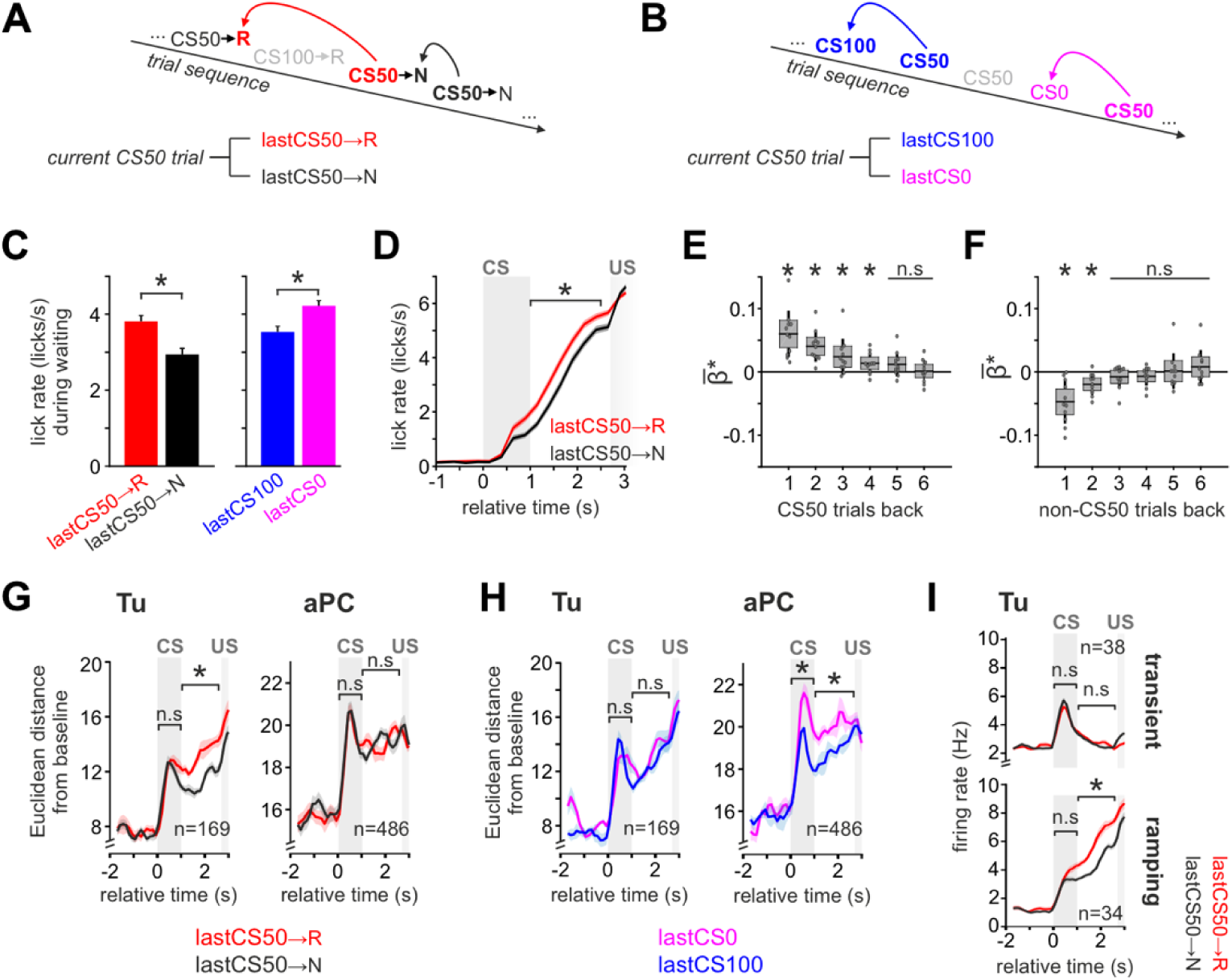
Reward prediction updating through the recent cue-specific outcome-history. (**A**) Scheme illustrating the division of CS50 trials based on their recent reward outcomes to test for dependence from outcome-history. CS50 trials were grouped according to the outcome of the previous CS50 trial being rewarded (lastCS50→R) or not (lastCS50→U). (**B**) To assess how the dynamic value updating in CS50 trials is influenced by reward outcomes paired to other stimuli, we separated CS50 trials depending on whether the preceding trial was CS0 or CS100. (**C**) Mean lick rate ± SEM during wait in CS50 trials was modulated by the outcome of the preceding trial (n = 88 in 11 animals; two-tailed paired t-test). Note that receiving a reward has a differential effect on the lick rate during the next trial depending on the stimulus identity. (**D**) Licking intensities differentiated CS50 trials throughout the waiting interval depending on the outcome of the previous CS50 trial (two-tailed paired t-test). Displayed average lick rates ± SEM. (**E**) Multiple Poisson regression with anticipatory licks in CS50 trials (during delay window) as dependent variable and the outcome of the previous six CS50 trials as independent variables (one-sample t-test on the resulting beta, n = 88 sessions in 11 animals). Anticipatory licking in CS50 trials was positively modulated by the outcome of the last four CS50 trials. (**F**) Same as (E) but with the outcome of the previous six CS100 and CS0 trials as independent variables. Anticipatory licks in CS50 trials were negatively modulated by the outcome of the prior non-CS50 trials. (**G**) Mean Euclidean distance from baseline ± SEM of the population vector during the lastCS50→R and lastCS50→N trials. If the previous CS50 trial had been rewarded, Tu population displayed a stronger response than if it had been unrewarded (left). In contrast, no recent outcome-history effect was observed in the aPC population (right). Two-tailed paired t-test at CS and during waiting. (**H**) Same as (G) but dividing CS50 trials based on whether the prior trial was CS100 or CS0. Satiety did not change the population response of Tu at CS and during wait (left), but reduced the population responses of aPC (right). Two-tailed paired t-test. (**I**) Recent outcome-history analysis for the transient- and ramping-clusters of Tu. Displayed: mean firing rate ± SEM during lastCS50→R and lastCS50→N trials for the two clusters. The ramping-clusters encoded outcome-history, while the transient-cluster showed no effect. See Fig. S7C-F for a systematic analysis of outcome-history and satiety in all response-clusters of Tu and aPC. In the figure: * indicates p < 0.05 (see Table S1 for exact p-values and test details) and n indicates the number of units.

We then tested whether the task-related global population activity would also reflect such outcome-history. Confirming the results from the analysis of the BOLD responses, we observed that the recent CS50 outcome-history was encoded by enhanced responses in the Tu (**Fig. 7G, left**), but not in the aPC (**Fig. 7G, right**). In contrast, satiety did not change the population response in Tu (**Fig. 7H, left**) but negatively modulated the response to CS50 in the aPC (**Fig. 7H, right;** see also **Fig. S7E-F**). While monotonic RP was encoded by both transient and ramping Tu populations (cf. Fig. 5), RP updates by the cue-specific outcome-history were significant only in the ramping population during waiting (**Fig. 7I;** see also **Fig. S7C-F**). Notably, in the ramping population, satiety had the opposite effect of the cue-specific history (**Fig. S7D**).

In summary, the ventral striatal circuit performed parallel computations of the RP in two neuronal populations. One population responded to CS and provided a stable representation of the RP. The second population encoded the anticipated RP during the waiting period before US and integrated the expectation update based on the recent cue-specific outcome-history. Cue-specific updating of the RP differed in its direction from satiety that modulated more prominently the aPC.

## DISCUSSION

### Mesoscale fMRI in the hierarchical approach

The hierarchical approach developed in this study may serve as a discovery strategy for key circuits in complex behavioral tasks and to reconcile findings in selected brain regions at the system level. The hierarchical approach starts from behavioral modeling to then identify task-relevant circuits in an unbiased fashion with functional imaging. Upon such regional identification, the cellular and network coding mechanisms are studied at finer grain resolution by means of electrophysiological recordings; thereby providing a systematic cross-scale integration from behavior, over mesoscale networks, to cellular functions.

Functional MRI in awake rodents offers the opportunity to assess the functional recruitment of deep and superficial circuits during task performance. Several aspects are critical when interpreting BOLD responses. Seminal work has provided insights into the neuronal basis of BOLD responses to isolated sensory stimuli in animals (Bartels et al., 2008, Logothetis et al., 2001, Nir et al., 2007). Even though BOLD changes reflect activity patterns of projection neurons, in some regions, such as the striatum, BOLD and rate changes can have an inverse relation (Mishra et al., 2011, Sloan et al., 2010). Moreover, little is known about the relation between population rate coding and BOLD signals during complex tasks. BOLD responses correlate with regional blood flow that is regulated for instance through nitric oxide synthesized by interneurons in an activity-dependent fashion (Cauli et al., 2004, Enager et al., 2009, Lee et al., 2020). These interneurons can, but do not need to, be positively correlated with principal neuronal activity (Cauli et al., 2004). It is, therefore, possible that inputs from sensory or reward coding regions may differentially innervate and drive these interneurons. These factors may eventually generate the observed opposing BOLD contrasts for the RP and PE, even though principal neurons responded to both with the same direction of firing rate changes. Therefore, when using fMRI as a localizer for task-related activity as in this study, both positive and negative BOLD associations are informative but, at present, cannot serve to predict the specific character of the underlying neuronal activity.

Even though the mouse hemodynamic response function is several fold faster than in primates, it will only partially separate sequential task events that occur typically in the range of seconds, and should, therefore, be corroborated by neurophysiology with millisecond temporal resolution. Further, fMRI has a relatively low signal-to-noise-ratio even at high field strength, which we compensated with repeated imaging and an optimized training scheme to generate sufficient cohort sizes. As an additional source of noise, movement is of particular concern especially in behavioral rodent fMRI. We employed a combination of denoising methods to minimize head-motion effects, but cannot fully exclude a residual impact of such artifacts (see Methods section ‘Functional MRI denoising’ for a detailed discussion). Notably, in the two brain regions that were also examined with electrophysiology, the task information contained in fMRI was confirmed in the neuronal population activity. Taken together, this study supports fMRI as a powerful tool in the localization of distinct subnetworks involved in complex cognitive functions at the system level.

### Forebrain circuits in olfactory reinforcement learning

Through the mesoscale fMRI approach, we identified different contributions to RP and PE among the primary olfactory cortices, subcortical circuits and higher brain regions. While forebrain regions broadly contributed to value-related information, the complete monotonic RP involved a more narrow network.

In the olfactory cortices, monotonic RP was represented primarily in the AON, and partially also in the pPC. AON provides massive cortical top-down inputs to the MOB, it may therefore not be surprising that MOB also represented monotonic RP. This MOB-AON loop dynamically updated value by the recent outcome-history. A stark functional segregation among olfactory cortices was found for the aPC, despite its reciprocal connection with the AON and the dense input from the MOB. As detailed by single-unit recordings, aPC encoded at CS primarily stimulus detection, but no monotonic RP. During the waiting period, however, task-inhibited monotonic RP responses emerged, that reflected in equally negative BOLD correlates in the aPC. At US, all primary olfactory cortices uniformly contributed to the value computation and little to the reward component of the PE, except for partial involvement of the AON.

Functional segregation was also observed in higher-order cortices. The mPFC, OFC, and anterior insular cortex all contributed to monotonic RP, but differed in their contribution to the PE. While the OFC correlated both to the value and reward components of the PE at US, the insula contributed mainly to its value component, and the mPFC did not correlate significantly to PE assessment in our task. Prefrontal and insular regions prominently project to the lateral striatum and NAc as well as the Tu (Gehrlach et al., 2020, Zhang et al., 2017). In the Tu, additional direct inputs from the olfactory cortices and the MOB converge (Gehrlach et al., 2020, Zhang et al., 2017). Tu and NAc represented all RP and PE aspects in the olfactory reinforcement learning task, including dynamic value updating. In line with anatomical input gradients from sensory and reward regions (Ikemoto et al., 2015), posterior parts of the Tu correlated with the reward component of the PE at US. Also, PE correlates localized in the lateral NAc shell, consistent with its contribution to positive reinforcement (Yang et al., 2018). Finally, the dorsal striatum similarly reflected all aspects of RP, but was dominated by the reward components of the PE at US.

Supporting the translatability of these fMRI findings in mice, conditioning in human fMRI studies recruited homologous prefrontal and ventral striatal regions involved across stimulus modalities (Daw et al., 2005, Ferenczi et al., 2016, Hare et al., 2008, Li et al., 2011, Niv et al., 2012, O’Doherty et al., 2004, O’Doherty et al., 2003) as well as modality-specific correlates in olfactory cortices or insular regions (Gottfried et al., 2002, Howard et al., 2016, Li et al., 2006, Zelano et al., 2007, Zelano et al., 2011). The largely non-overlapping forebrain representations of reward and value components of the PE at US match their distributed computations in humans (Hare et al., 2008, Niv et al., 2012). Consistent with humans (Hare et al., 2008, Niv et al., 2012), the mouse NAc represented both reward and value of the PE at US. Other brain regions that also represented both PE components at US, specifically Tu and the MOB-AON loop, have not been explicitly examined in human whole-brain fMRI due to their comparatively small volume in humans in contrast to macrosmatic species. Among the densely interconnected network revealed by mouse fMRI, the Tu is uniquely positioned at the junction between the olfactory and reward system and sticks out because of its contribution to multiple facets of stimulus-outcome learning.

### Non-redundant parallel coding of reward prediction

Recent studies have revealed that Tu neurons are recruited by stimuli that predict rewards upon conditioning (Gadziola et al., 2020, Millman and Murthy, 2020, Murata et al., 2015, Oettl et al., 2020, Zhang et al., 2017). In these studies, the duration of the conditioned stimulus partially overlaps with US. To better capture reward anticipation, in the present study we separated the CS and US by a waiting gap. Upon learning, we observed the evolution of two prominent task-excited populations of Tu units computing either a transient stimulus-bound RP or a ramping anticipatory RP during the waiting period.

The transient stimulus-bound RP signal was encoded by the sum of distributed odor responses of multiple units, many of which were selective for only one odor, but that collectively encoded monotonic RP. This RP signal emerged at CS after cross-session learning and was little influenced by the recent outcome-history or satiety; consequently generating a stabilized representation of the RP. The stability of the RP representation at CS may relate to its formation through synaptic plasticity (Wieland et al., 2015). In fact, the stimulus response potentiation evolves gradually upon repeated pairing of the odor with phasic dopamine in awake mice (Oettl et al., 2020). Once emerged, these RP-coding CS signals in the Tu drive downstream the equivalent RP responses of VTA DAN (Oettl et al., 2020).

In contrast to this CS-bound signal, the ramping anticipatory RP signal evolved during the waiting interval after the stimulus has ceased. Ventral striatal anticipatory ramping activity is also found during waiting in head-fixed primates (Schultz et al., 1992) and thought to reflect anticipatory timing to reward (Gershman and Uchida, 2019, Mello et al., 2015) informing prefrontal cortices (Meck et al., 2008). Within the Tu, the ramping RP signal was encoded redundantly, with multiple single-units responding to all rewarded odors proportionally to their associated value. Value update was fast and stimulus-specific, in line with the modeled RP. The faster value updating of the ramping population could be interpreted as a neuronal implementation of the TD model, where values more proximal to the PE at US are more quickly updated than the distal ones at CS. While stimulus-bound RP responses drive DAN at CS, this is not the case for Tu ramping activity. In fact, DAN do not express anticipatory ramping in head-fixed trace conditioning (Cohen et al., 2012, Kim et al., 2020) but only when animals approach rewards in space (Howe et al., 2013, Kim et al., 2020, Mohebi et al., 2019). Finally, the enhanced responsiveness to reward of the anticipatory ramping population compared to the CS-bound RP population adds to their differentiation. Potentially, such differentiations could map on the anterior-posterior gradients for PE value and reward representations found here, different olfactory and reward-related projection gradients (Ikemoto et al., 2015, Wesson, 2020), or direct- and indirect-pathway Tu neurons (Murata et al., 2015). Taken together, the different coding strategies, the non-overlapping neuronal clusters, the functional dissociation of the two RP signals and VTA, and the different responsiveness to reward, support the presence of parallel networks for stabilized and dynamic reward prediction in Tu.

The hierarchical approach presented here, with its cross-scale analysis, can be applied broadly to reveal the computations in distributed brain networks and can foster the discovery of key brain regions and mechanisms in cognitive and translational sciences. Here, the approach identified the olfactory tubercle of the ventral striatum as one of the key circuits to compute multiple non-redundant olfactory reward predictions in parallel networks of projection neurons. While the CS-bound stabilized RP signal provides an initial safe estimate to drive dopamine midbrain neuron coding, the subsequent anticipatory RP dynamically integrates recent experiences in preparation of reward retrieval. This olfactory prediction coding hub operates within a network identified by fMRI of functionally-segregated olfactory and higher-order regions.

## ACKNOWLEDGMENTS

The authors thank Dr. Peter Dayan for discussions and comments on the manuscript and Drs. Urs Braun and Heike Tost for initial discussions. We thank Ariana Froemmig, Felix Hoerner and Catrin Loeb for excellent technical assistance. This work is supported by the BMBF grant n. 01GQ1708 to W.K., Boehringer Ingelheim Foundation grant ‘Complex Systems’ to W.K., DFG Priority Program SPP1665 KE1661/2-2 to W.K, the Ch. and H. Schaller Foundation to E.R., and the Clinician Scientist Program “Interfaces and Interventions in Complex Chronic Conditions” by the German Research Foundation (DFG) (EB187/8-1) to J.R.

## AUTHOR CONTRIBUTIONS

C.C.v.H., W.K., A.M.L. and L.W. conceived the project. M.Sch. designed the recording array and behavioral setups. L.W. performed behavior, fMRI, and neurophysiology. C.F. designed and implemented the behavioral model. M.Sa., R.B. and L.W. developed the pupil analyses script. W.K., E.R. and L.W. designed and implemented the analysis for the neurophysiology. D.W. helped with the neurophysiology. M.F.G., C.C.v.H., R.H. and L.W. designed and implemented the fMRI analyses. A.S., J.R. and W.W.F. provided scripts, resources, and helped to implement the fMRI analyses. C.C.v.H., W.K., E.R. and L.W. wrote the manuscript.

## DECLARATION OF INTERESTS

The authors declare no competing interests.

## MAIN TABLES

**Table S1.**
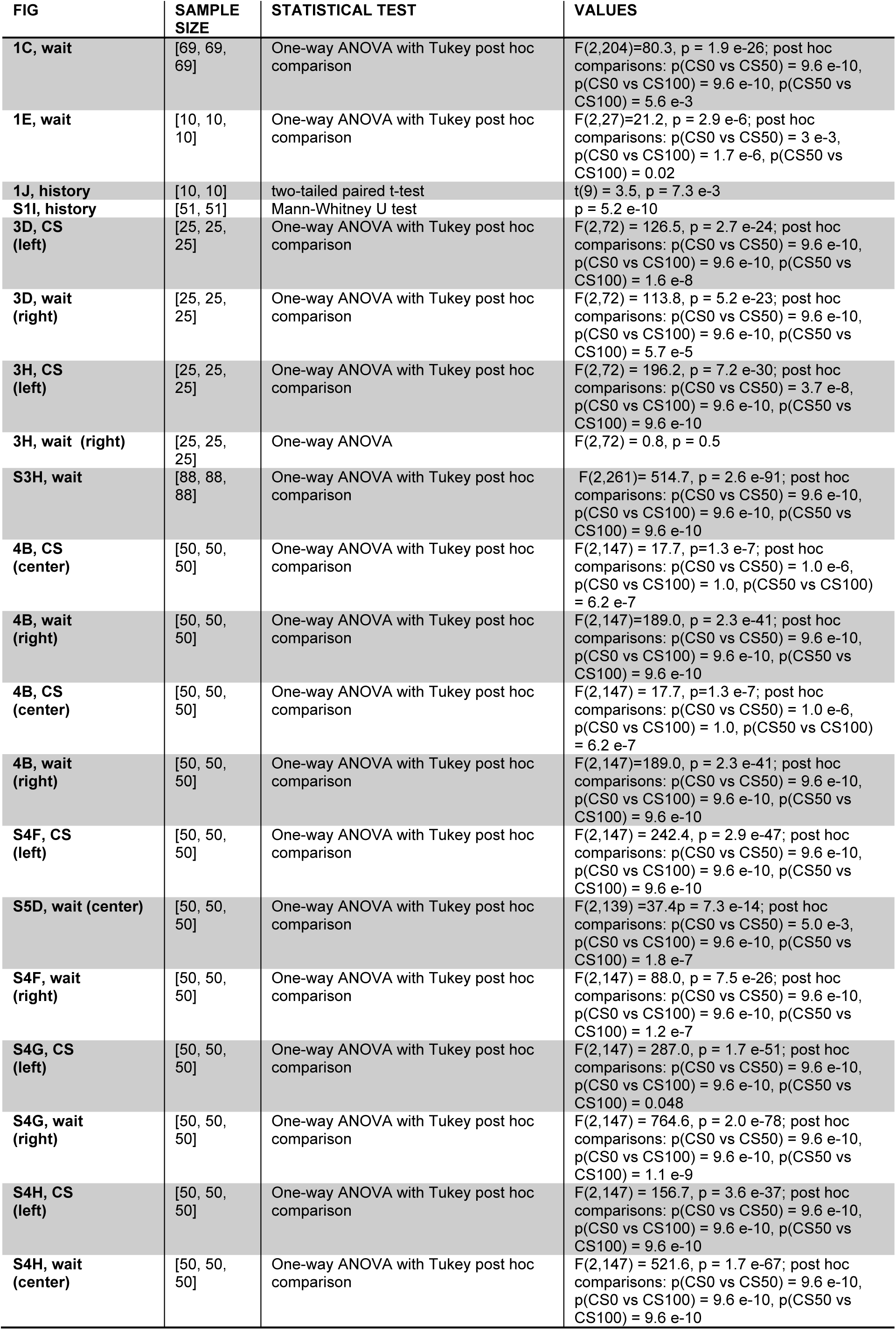
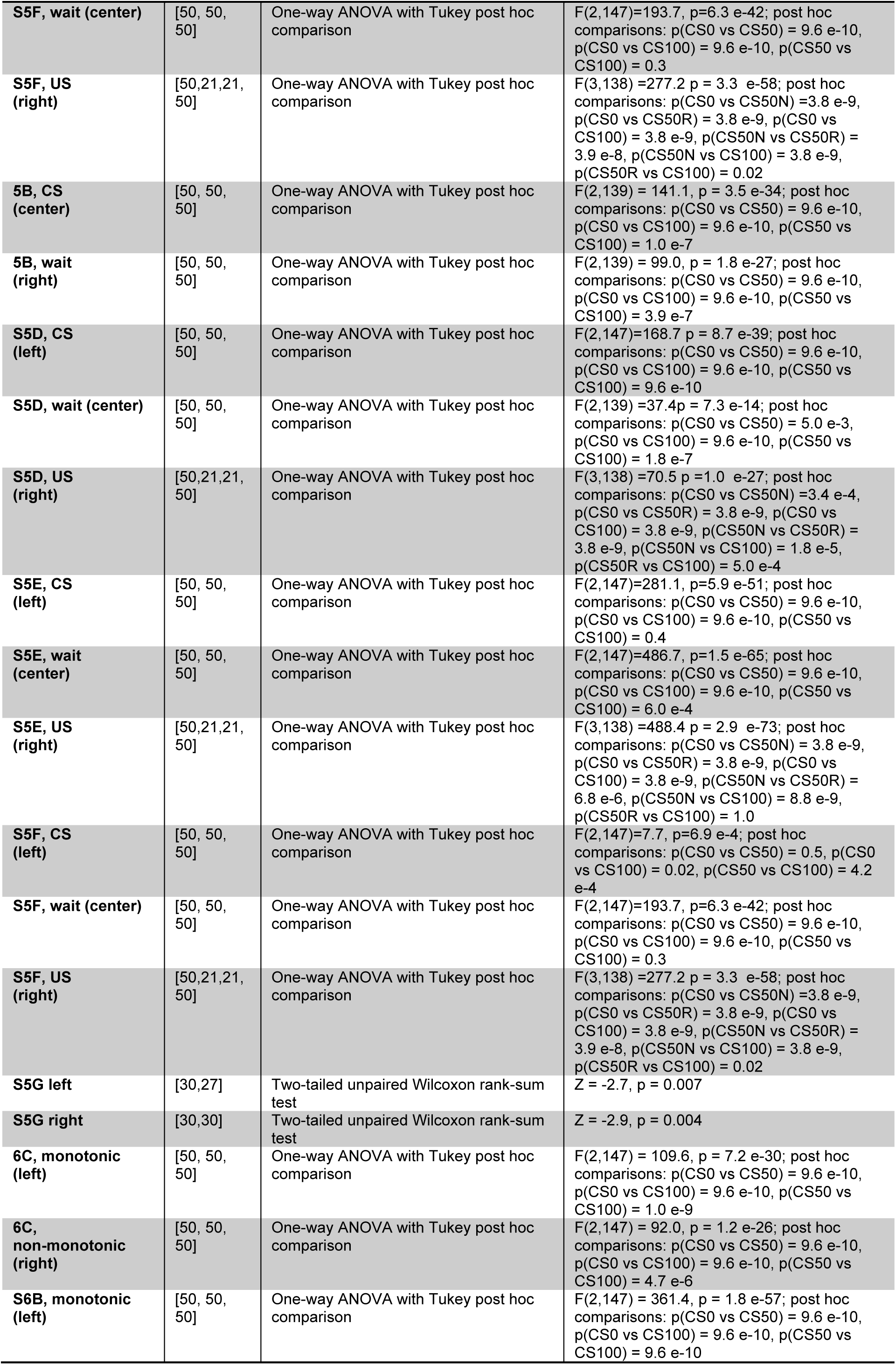
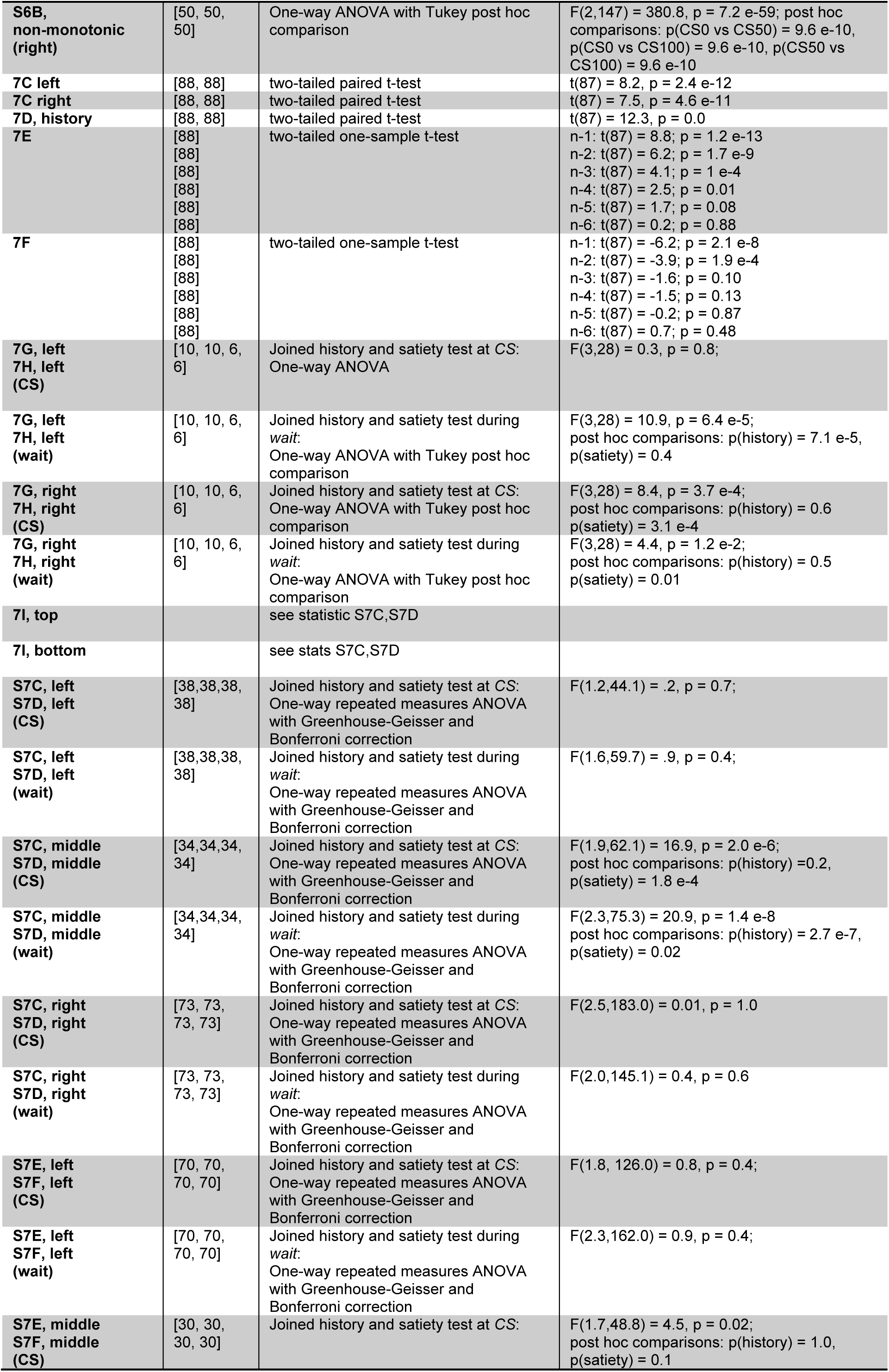
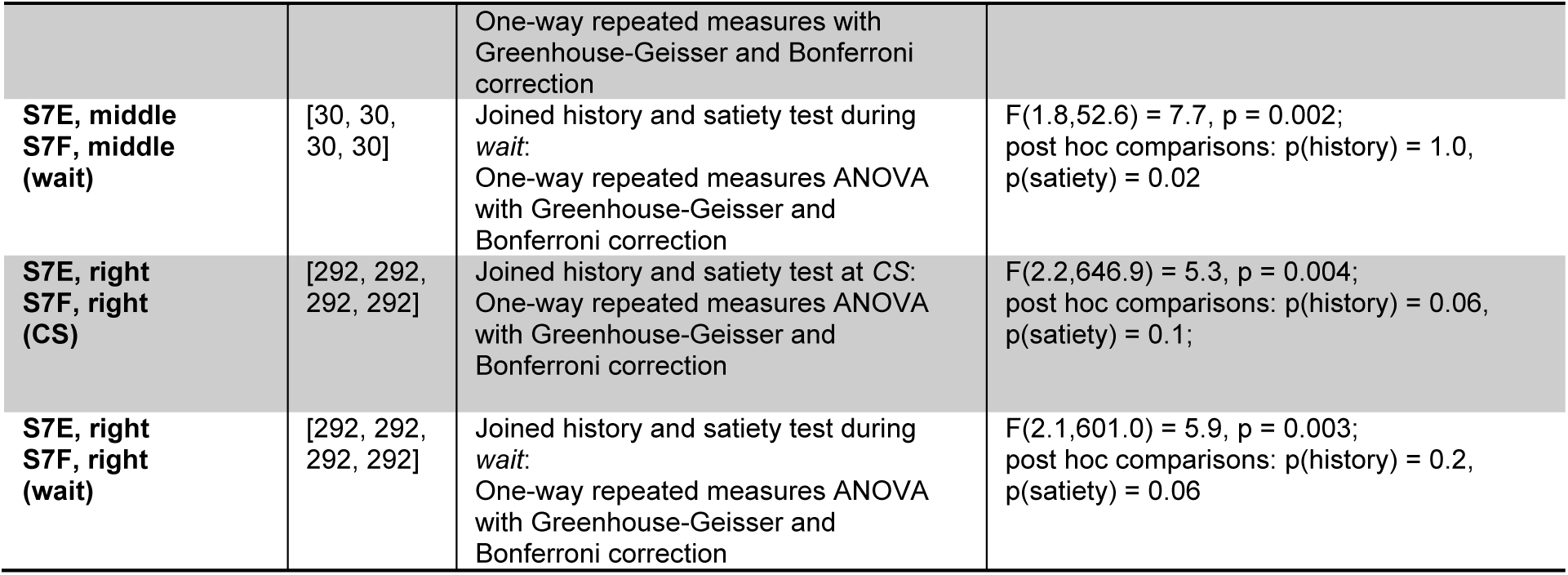
Statistical tests. Description and results of the statistical tests displayed in the manuscript figures. The first column of the table indicates the figure number and panel of the corresponding figure.

## METHODS

### RESOURCE AVAILABILITY

#### Lead Contact

Further information and requests for resources should be directed to and will be fulfilled by the Lead Contact, Dr. Wolfgang Kelsch.

#### Materials Availability

This study did not generate any new animal models nor reagents.

#### Data and Code Availability

At the time of publishing, the behavioral, modeling, fMRI, neurophysiological data will be available upon reasonable request from the corresponding author.

### EXPERIMENTAL MODEL AND SUBJECT DETAILS

#### Animals and Husbandry

Male C57BL/6N mice were obtained directly from Charles River Laboratories (23 animals for the fMRI measurements, 11 animals for single-unit recordings). Mice were housed individually in a standard 12 hours light-dark-cycle. Food and water were given ad libitum, except when water supply was controlled for behavioral training. All procedures were in accordance with the National Institutes of Health Guide for the Care and Use of Laboratory Animals and the EU 2010/63 directive, and approved by the local animal welfare authority (Regierungspräsidium Karlsruhe).

### METHOD DETAILS

#### Behavioral Modeling and functional MRI

##### Implantation of the head bar

All surgeries used standard aseptic procedures and conformed to common veterinary practice. Analgesia (Metacam, Boehringer Ingelheim) was administered before and after surgery. At least 12-week-old mice were anesthetized with isoflurane. The animals were then transferred to a stereotactic apparatus with non-rupturing ear bars and placed in a custom-built platform of the same dimension as the MRI cradle. It was ensured that the fixed licking spout of the MRI cradle was at the same height as the animal’s lower lip at a distance of 5 mm. A roughly circular flap of skin was removed from the skull and local anesthesia was administered. The lateral and nuchal muscle insertions were left intact. The skull was cleaned and disinfected. Tissue adhesive (3M Vetbond) was applied to the margins of the skin attached to the circumference of the exposed skull to avoid soft tissue damage and contamination. The remaining periost on the exposed skull was removed. A layer of dental glue (C&B Superbond, Sun medical) was applied on the (inter)parietal bone, followed by a layer of dental cement (Kulzer Palladur) that connected the custom-designed head bar produced with stereolithography (Accura55, 3D-Systems).

##### MRI-compatible behavioral setup

We developed an MRI-compatible setup for the odor-guided reward learning task (see **Fig. 2, S1, S2**). The apparatus comprised of an MRI cradle, an olfactometer, a programmable syringe pump for reward delivery (AL4000-220, World Precision Instruments), an optical licking detector, and Arduino microcontrollers (Arduino Mega 2560). Odors were delivered using a custom-made olfactometer. Odors were kept in liquid phase (diluted 1:100 in mineral oil) in dark vials and mixed into a nitrogen stream that was further diluted by 1:10 into a constant air stream in the olfactometer. The following natural flower odors were used: geranium, ylang-ylang and rose (Sigma Aldrich W250813, W311936 and W523704, respectively). Water as reward was delivered through an independent tubing system and controlled by a high-precision water pump. Odorized air and water were both guided from the setup located in the control room to the animal bed through 1/32 inch I.D. inert PTFE tubing (NResearch) connected to the odor and lick ports. Considering the 3 m long tubing used to deliver the odorized air to the odor port, the steepness of the odor onset (20 ms to max) and its latency after final valve opening (400 ms) was regularly controlled by a photo-ionization detector (miniPID, AuroraScientific). Odor valves and syringe pump were controlled through Arduino microcontrollers. Licking behavior was measured using a custom-made MRI-compatible optical lickometer. Infrared light was delivered via fiber optics and miniature roof prisms from an LED source (Thorlabs M660F1; 660nm), collimated at the lick port with lenses (Thorlabs 354140-B) and returned to the control room via another optic fiber to the detector (Thorlabs DET10A2). Animals were positioned so that their tongue broke the beam on each lick. Outputs from the olfactometer, the optical lick detector, the water pump and TTL pulses from the scanner were recorded with the same RHD2000 interface board (Intan Technologies) with a sampling frequency of 1 kHz. For training outside the scanner, a custom-made simplified MRI cradle was used.

##### Behavioral training

Mice were trained in a trace conditioning task to learn associations of stimulus-outcome pairs. Three days before behavioral training, water-intake was controlled in their home cages (90% of baseline body weight was targeted). Body weight was monitored daily and always maintained above 85% of baseline body weight. Mice were placed in the head-fixed setup for habituation. The habituation sessions did not exceed 15 minutes. In general, mice habituated to head fixation after 2-3 days. Then, conditioning sessions started. Each trial started with 1 s of odor presentation followed by a waiting period of 1.7 s. Reward (5 μl water) was delivered immediately after the waiting window (see **Fig. 1A, middle**). Reward timing and reward size were not varied. Licking responses had no influence on whether reward was delivered or not. The inter-trial interval was randomly drawn from a uniform distribution between 10 and 12 s. The training comprised of two stages. In *Stage 1, a* single odor was presented and rewarded at 100%. When mice licked consistently, they progressed to the next stage. *Stage 2* corresponded to the final paradigm and consisted of three distinct odors delivered pseudorandomly to keep the proportion of odors constant between sessions. No stimulus was consecutively applied more than three times in a row. No more than three consecutive trials were rewarded. Animals performed 150 trials per session (in the MR-environment, 20-30 trials of which were regarded for acclimatization). Odor cues predicted reward at 100%, 50% and 0% (hereafter, the odor cues were labeled CS100, CS50 and CS0, respectively) (see **Fig. 1A**). Across sessions, the odor and reward contingency pairs were unchanged. The first session of Stage 2 was considered the ‘first training session’ in **Fig. 4 and 5**. Generally, CS100 and CS50 were treated as ‘Go’ cues and odor CS0 as a ‘No-go’ cue. If mice licked at least three times during the anticipatory window (from 1.5 s to 2.8 s after odor onset) or in the reward window (from 2.8 s to 4.1 s), it was considered as a go-response. Fulfilling the lick criterion during ‘Go’ trials was regarded as ‘Hit’, while it was assessed as ‘False alarm’ during ‘No-go’ trials. Not meeting the lick criterion was regarded as ‘Correct rejection’ for ‘No-go’ trials and as ‘Miss’ for ‘Go’ trials. There was no punishment for false choices. The performance within each session was evaluated by calculating the correct rate’:

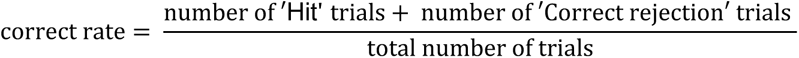

After mice designated for functional imaging reliably performed the task above criterion (> 80% correct trials, normally after 6 to 12 training sessions), a sham coil was placed above the head and recorded MRI pulse sequence noise was replayed in ‘mock scanning sessions’ to acclimate mice to the MRI environment. Sound levels in the mock scanner were increased gradually to the noise levels in the scanner bore. In the behavioral analysis, only sessions with a ‘correct rate’ higher than 80% were included.

##### Pupil imaging

Pupils were imaged unilaterally in trained mice (n = 10 sessions with one session per mouse). Pupil data was collected in the recording chamber with ambient light illumination (blue LED, 465 nm). To fully capture the pupil dynamics during task performance, the intensity of the LED light was set so that the pupil was moderately dilated. DinoCapture 2.0 software was used as video recording system. Videos were acquired at 20 frames per second (1280 x 1024 pixel) with a digital infrared USB camera (AD4113T-I2V Dino-Lite Pro2 digital microscope) providing infrared illumination by LEDs (940 nm). Infrared illumination did not affect pupil diameter. At the beginning and at the end of each session, an additional infrared LED was switched on for 1 s in order to generate timestamps. The timestamps were then used for post-hoc alignment of pupil and behavioral data.

##### Pupil diameter analysis

Pupil diameter was detected frame-wise using in-house developed MATLAB scripts based on a pulse-coupled neural network (PCNN) algorithm. Briefly, pupil videos were read into MATLAB and converted to grayscale. Then, pupil videos were automatically centered and cropped around the pupil. Based on the adjusted frames, the PCNN algorithm calculated binary images reliably segmenting the pupil from the surrounding tissues. A convex hull was fitted to the contour of the detected pupil area. The pupil radius was detected for each frame by calculating the mean distance between all points on the convex hull and its midpoint. Outliers in the detection, e.g. during blinks, were excluded from the data by removing the entire trial (exclusion criterion: increase or decrease in pupil diameter of more than 5% within 2 frames). Accuracy of the algorithmic fit was verified visually for each session. Pupil diameter was expressed as a percent change from baseline. Baseline was derived from the average pupil diameter in the time window from −2 s to 0 s relative to odor onset. Mean responses were calculated for each trial types for each animal and, in turn, averaged to get the group response ± SEM. Pupil data were tested for normality using a Kolmogorov-Smirnov test. An ANOVA test was performed between CS100, CS50 and CS0 trials during the waiting window (from 1 s to 2.5 s relative to odor onset).

##### Reinforcement learning model

In probabilistic conditioning tasks, animals attempt to predict future outcomes. This reward prediction (RP) signal is adjusted in time through the difference between the predicted and the actual outcome, namely the reward prediction error (PE) at US. In order to parametrize the PE and the RP, we modeled a temporal difference model TD(0) on a trial basis, with no eligibility trace or discount factor (O’Doherty et al., 2003, Seymour et al., 2004).

For each trial, we considered three time points: t = 0 for the baseline before the trial started, t = 1 for the odor presentation (CS), and t = 2 for the reward delivery (US). The states at time t = 1 and t = 2 were defined by the stimulus presented at that time (CS0, CS50 or CS100), each of them having a corresponding value *V*(s) and an outcome, *r*, representing the reward, written as dichotomous variable {1,0} (1 for reward, 0 for no reward). At time t = 0, *V*(0) represented the prediction before the trial started. In the TD(0) model, the PE reflects the internal computation of the difference between two successive predictions. Thus, at each time point, t, the PE is defined as:

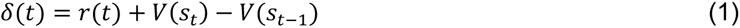

Note that, at the reward time point, we assumed *V*(*s_t_*) = 0. This is because learning at US is quick, and the animals, rather than learning an internal value, react directly to the reward. Thus, at US, the PE was computed as: *δ*(*US*) = *r* - *V*(*s*_*t*-1_), that is the difference between the predicted and the actual outcome at US. At CS time, in contrast, where no reward was delivered, we had *r* = 0, thus the PE was: *δ*(*CS*) = *V*(*s_t_*)-*V*(*s*_*t*-1_). The value of *δ*(*t* = 0) before the trial start was 0. The state expectation values were updated on a trial-by-trial basis through *δ* according to the TD learning rule:

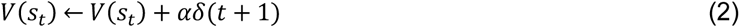

where *α* is the learning rate. Our experiments were not designed to disentangle experimentally *V* from *δ*(*CS*) (Kim et al., 2020).

##### Parameter estimation and selection

The learning rate *α* was set as free parameter of the model and estimated using pupillary data from the pupil imaging sessions described above (n=10 sessions). The average change in pupil dilation *d* was modeled as function of the expected value *V* and the PE *δ*:

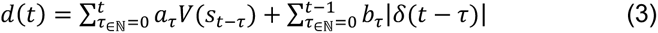

where *t* = {0, 1, 2} (0 for the baseline starting point, 1 for the odor presentation and 2 for the reward time). *a_τ_* and *b_τ_* were set as free model parameters. *d*(*t*) indicated the average change in pupil dilatation between two successive times in the trial: we defined d(0) as the average change in dilation from before the trial start to the odor presentation; then d(1) as the change from the odor presentation to the reward delivery, and d(2) as the change after reward delivery. The interval before the trial started, namely the interval between t = 0 and t = 1, was defined from −2 s to 0 s relative to odor onset in line with the pupillary analysis (see above). It was selected long enough to yield a stable average. The interval from t = 1 to t = 2 was defined from 0 to 2.7 s, and, for consistency, the interval after US was chosen to be from 2.7 to 5.4 s. The model parameters *ϑ* = {*α*, *α_τ_*, *b_τ_*} were estimated minimizing the distance between the modeled (eq. 3) and the real average change in pupil dilation.

After the average learning rate *α* had been estimated on the basis of the 10 sessions with pupil imaging, we used this *α* to build a temporal difference model for 151 different trial realizations. Of those, 100 realizations were simulated on as many CS-US sequences, randomly generated with the algorithm used for the behavioral sessions, and used to produce Fig. 1F,I; and 51 were simulated on the actual trial sequences of the fMRI sessions and used in the fMRI analyses. We used the learning rules in eq. 2 with the PE defined in eq. (1) (cf. Fig. 1F,G) to model the learning. In this way, we formulated specific predictions on RP updates and regressed them on fMRI data of mice performing the task in the scanner to identify key forebrain regions. The initial condition of the values of all stimuli was set to 0 in the simulations of Fig. 1, and set to 0, 0.5, and 1 for CS0, CS50 and CS100 respectively in the fMRI sessions (as the animal were already performing when the fMRI scanning started, c.f. Fig S1F).

##### Functional MRI acquisition

Experiments were conducted on a small-animal 9.4 Tesla MRI scanner (94/20 Bruker Biospec, Ettlingen, Germany) with Avance III hardware, BGA12S gradient system with the maximum strength of 705 mT/m and running ParaVision 6 software. We utilized a whole-body linear volume transmitter coil combined with an anatomically shaped 4-channel receive-only coil array. Mice underwent fMRI scans after habituation to the MRI environment (see above). Awake mice were head-fixed in the MRI cradle: their trunks were gently held within a plastic cylindrical tube to further prevent massive body motion. Once positioned inside the scanner bore, the task was initiated, allowing the subject to orient to the task and minimize distress. The 20-30 trials presented before onset of the fMRI sequence were not recorded. A localizer sequence was used to confirm the correct positioning of the animal and a field map was acquired. Before the fMRI scan started, the main magnetic field (B0) was homogenized by automatic shimming. The fMRI time series were acquired while mice performed the odor task using an echo-planar imaging (EPI) sequence with the following parameters: TR/TE: 1300/17 ms; flip angle: 50°; 21 slices; matrix size: 64 x 64; slice thickness: 0.5 mm; interslice gap: 0.1 mm; voxel size: 0.25 x 0.25 x 0.6 mm; 1400 volume acquisitions. Temporal resolution was increased by restricting the field-of-view to rostral parts of the brain, containing the olfactory, frontal and striatal regions. Each EPI session lasted 24 minutes. The behavioral session was followed by a high-resolution T2-weighted RARE anatomical image acquisition (TR/TE 1200/50 ms, matrix size 96 x 113 x 48, voxel size 0.16 x 0.16 x 0.31 mm, RARE factor 16). To increase statistical power, animals were repeatedly measured (up to 4 sessions per animal), but not more than once per day. Mice that did not lick at reward in the scanner were excluded from further measurements to avoid unnecessary distress. This yielded a dataset of 67 sessions from 23 animals. Of these, 51 sessions from 18 animals fulfilled the performance criterion described above and were included in fMRI group statistics.

##### Image data processing

All data were processed using Statistical Parametric Mapping version 12 (SPM12) (http://www.fil.ion.ucl.ac.uk/spm/) and custom-written MATLAB scripts. fMRI data were preprocessed with the following steps (Clemm von Hohenberg et al., 2018): discarding the first five volumes in the series to avoid influences of magnetization before the scanner achieves steady state, correction for head movement by realignment to the middle volume, correction for geometrical distortions using the acquired field maps, slice-timing correction, and spatial normalization to a mouse brain template in the Paxinos stereotactic coordinate system, by applying the non-linear normalization parameters of the structural images to the functional images. Normalized functional images were additionally smoothed with a 0.6 mm isotropic Gaussian kernel.

##### Functional MRI denoising

Functional MRI is susceptible to movement artifacts, which is of particular concern given the association between the paradigm, licking activity and corresponding head motion. To quantify head motion, absolute values of the differentiated realignment parameters’ first derivative were averaged over all sessions for the three translations and three rotations (see **Fig. S2B**). To minimize motion artifacts, we explored several denoising methods. Regression of realignment parameters, which is widely applied, has several shortcomings, as it depends on the accuracy of the realignment process, does not reliably capture non-linear or delayed effects, is anatomically unspecific and does not differentiate between task-correlated neuronally-based activity and artifacts (Caballero-Gaudes and Reynolds, 2017, Pruim et al., 2015). The latter effect is of particular concern in the context of task-based fMRI, where motion is typically correlated with the effect of interest. Indeed, we found that plausible gray-matter activation in response to rewarded CS types (e.g., in dorsal striatum) was substantially decreased (while areas of deactivation became more extended) when the six realignment parameters and their derivatives were included in the GLM, exclusively or in addition to the other denoising methods described subsequently.

Removal of high-motion volumes (termed “scrubbing” or “censoring”) has been shown to outperform motion regression under certain conditions (Power et al., 2014, Siegel et al., 2014), but again suffers from important drawbacks, namely that it is difficult to find valid criteria for frame removal, especially in the context of task-based fMRI (Siegel et al., 2014). Further, excessive amounts of data may be removed with corresponding loss of temporal degrees of freedom (Patel et al., 2014, Pruim et al., 2015), even when delayed movement effects are not addressed. In task-based fMRI, censoring may inherently introduce bias and disproportionately affect image frames of certain kinds, such as during rewarded trials or in high motivational states.

Other approaches have been proposed that allow for a more specific removal of presumed motion artifacts, by considering their typical spatial and/or temporal characteristics. Among these methods, we explored Wavelet Despiking, which exploits the divergent frequency characteristics of motion artifacts, and despikes the BOLD timecourse *locally* in a temporal, spatial and frequency sense (Patel et al., 2014). However, Wavelet Despiking has not been systematically assessed in task-related fMRI, where the frequency spectrum may be more heterogeneous than in resting-state fMRI. Additionally, in rodents, the hemodynamic response function (HRF) is faster than in primates (Chen et al., 2020, Lebhardt et al., 2016, Schlegel et al., 2015) and therefore has a weaker lowpass filtering effect, potentially allowing true neuronally-based BOLD signal to contain higher frequencies. In line with this, we found Wavelet Despiking to introduce implausible temporal “smearing” such that activations started slightly before the trial. This would have been of particular concern since events were relatively close in time in our paradigm. Yet, the anatomical patterns of BOLD responses were not substantially altered by Wavelet Despiking in our data (data not shown).

We therefore, alternatively, employed group-independent component analysis (ICA) for removing motion-related noise, using the fastICA toolbox (http://research.ics.aalto.fi/ica/fastica/) (Han et al., 2019). ICA-based denoising exploits typical anatomical patterns of motion artifacts, and has been applied in previous awake rodent imaging (Han et al., 2019, Tsurugizawa et al., 2020). In our group-ICA, four (out of 25) components were manually identified as motion-related (see **Fig. S2C**); their session-specific timecourses were removed before reconstructing the images. Criteria for component removal were typical location at the edges of the brain and/or in and around ventricles. One widespread, anatomically unspecific component (C1 in Fig. S2C) was also removed due to high correlations with realignment parameters and with the mean CSF timecourse. ICA was performed after other preprocessing including smoothing, to maximize signal-to-noise ratio and anatomical overlap between sessions/subjects for ICA, and also in accordance with previously described ICA denoising procedures (Han et al., 2019, Pruim et al., 2015). As described below, we included cerebrospinal fluid (CSF) timecourses in the SPM general linear model, to further account for physiological and non-physiological noise. Lick events were also included in the model (see below), which served as an additional control for the effects of head motion, which is highly correlated with licking activity.

##### Functional MRI analysis

As mentioned, the HRF varies significantly between species, with faster kinetics in mice compared to humans (Chen et al., 2020, Lebhardt et al., 2016, Schlegel et al., 2015). We therefore used a mouse-specific HRF estimated by Lebhardt et al., 2016. In all models, CS and US timepoints were modelled as events (“stick functions”, duration 0 s).

Specifically, we computed three different general linear models (GLM; SPM12) at the session-wise analysis level:

First, to locate the brain regions encoding reward prediction, we modelled all CS timepoints (irrespective of trial type) as one event type, and parametrically modulated this with the *V*(*CS*) estimated from the TD model. The four event types at the US timepoint (reward after CS100; reward after CS50, non-reward after CS50, non-reward after CS0) were each modelled with a separate event regressor. In this model as in all other models, lick events were also modelled as events, to disentangle neuronal correlates of motor activity from correlates of reward expectation and PE *per se*. Additionally, time series averaged over cerebrospinal fluid voxels were added as a nuisance regressor.

Secondly, to disentangle more specific components of value coding, especially monotonic RP and recent outcome history, we created a separate GLM, where all CS event types (CS100, CS50 and CS0) and all US event types (reward after CS100; reward after CS50, non-reward after CS50, non-reward after CS0) were each modelled with a separate stick function regressor. The CS50 regressor was parametrically modulated by the recent CS50 outcome history (namely whether the last CS50 trial had been rewarded or not). Analyses of monotonic RP and outcome-history were restricted to those voxels that were significantly associated with *V*(*CS*) at the group-level, based on the first GLM.

Thirdly, to test which brain regions encode the PE components (–*V*(*CS*), *r*), we computed a GLM where the three CS event types (CS100, CS50, CS0) were modelled with separate stick regressors, and all US event types were modelled by one single regressor, and the latter was parametrically modulated by –*V*(*CS*) and *r*. Of note, we did not orthogonalize one of the two parametric modulators with respect to the other, thus ensuring that only the unique variability of each parametric modulator is attributed to it.

Statistical maps representing the betas of the regressors of interest were then fed into group statistics. For this, we used the SPM12-based Sandwich Estimator toolbox (SwE). SwE is specifically designed for repeated-measures neuroimaging data and allows the number of sessions per subject to vary (Guillaume et al., 2014). The equivalent of one-sample t-tests was used within the SwE framework to test for effects on the group level. Analysis was restricted to gray matter. Unless otherwise stated, statistical threshold was set to p < 0.025 false discovery rate (FDR)-corrected, for two-sided testing.

#### Electrophysiological Recordings

##### Recording array

We used an in-house designed tetrode array for dual-site recordings described in detail in Oettl et al., 2020. In brief, custom-designed printed circuit boards (PCB) (±10 μm, Würth Electronics) with a soldered Molex SlimStack connector served as an electrode interface board (EIB). The tetrodes, spun from 12.5 μm teflon-coated tungsten wire (California Fine Wire), were placed parallel to each other with the help of a guiding scaffold. The tetrodes were fixed to the PCB with a drop of liquid acrylic adhesive. For electrical contact, the single wires were soldered to the EIB after threading them through 200 μm vias and through-holes. For protection, the single wires were coated with two-component epoxy. The tetrode tips were gold-plated with a NanoZ-device (Multi Channel Systems) targeting an impedance of 300 kOhm. During recordings, the arrays were connected to the IntanRHD2164 head stages using a custom-built adapter (Molex SlimStack connector to two 36 Omnetics Nano Strip connectors). A custom-built titanium head bar was glued to the array.

##### Implantation of recording array

Eleven male C57BL/6 mice were implanted with two 16-tetrode arrays, each positioned in Tu and aPC. Pre- and post-surgery analgesia was administered. Mice were anesthetized with isoflurane and attached to a stereotactic apparatus with non-rupturing ear bars. A roughly circular flap of skin was removed from the skull and local anesthesia was administered. The lateral and nuchal muscle insertions were left intact. Tissue adhesive (3M Vetbond) was applied to the margins of the remaining skin and holes drilled into the skull above the brain regions-of-interest and above the cerebellum for grounding. We then coated the skull using Super-Bond C&B (Sun Medical). An insulated copper wire attached to the recording array was connected to a small gold pin (Neuralynx) and placed into cerebellum. The tetrodes were inserted into the target regions with a motorized 3-axis micromanipulator (Luigs&Neumann) (target coordinates relative to the center of the respective array for Tu: 1.6 mm anterior to bregma, 1.3 mm lateral from bregma and 4.9 mm ventral from dorsal brain surface; target coordinates for APC: 1.4 mm anterior to bregma, 2.3 mm lateral from bregma and 3.8 mm ventral from the medial dorsal brain surface). Once the target depth was reached, dental cement (Kulzer Palladur) was applied to fix the recording array in the final position. Animals recovered in their home cage and were monitored after surgery.

At the end of recording sessions, mice were euthanized and perfused with paraformaldehyde (4%). Due to low levels of scar formation and microglia activation with this tetrode array, we could not reliably detect fiber tracts in histological sections with Nissl staining. Therefore, the head was postfixed in PFA 4% for at least 2 weeks before sectioning. Rostrocaudal and mediolateral placement in the borders of Tu and aPC were confirmed histologically before sectioning (see **Fig. S3B**). Subsequent removal of the tetrode array left visible scars in the tissue so that the recording location could be confirmed by post-hoc histological examination (see **Fig. S3C**).

##### Electrophysiology head-fixed set-up

The same olfactometer and behavioral control system setup configuration as in the fMRI was used for the electrophysiological study. Single-unit recordings were conducted using two Intan 64 channel RHD 2164 miniature amplifier boards connected to the RHD2000 interface board. Software provided by Intan Technologies was used for data storage. Data were sampled at 30 kHz during neural recordings.

##### Behavioral paradigm

The same training procedure, task, performance criteria, and behavioral analyses were applied to the electrophysiology cohort as for the fMRI cohort.

##### Multiple Poisson regression on the licking data

To assess whether the anticipatory licking in the delay period of CS50 trials was influenced by (1) the cue-specific outcome of the previous CS50 trials (outcome-history) and by (2) the outcome of previous CS100 and CS0 trials (satiety), we conducted two separate multiple Poisson regressions on the licking data (see **Fig. 7E-F**). The Poisson model was defined by the equation: log(*μ_t_*) = *β* · *X* where *μ_t_* is the expected value of the anticipatory licking in the current CS50 trial, *X* is the regressor matrix and the *β* are the regression coefficients. The columns of the matrix *X* are the regressor vectors. The elements of the regressor matrix assumed values *X_ij_ ∊* {1, −1}, where 1 codes for a rewarded trial and −1 for an unrewarded trial. In case (1), the regressor vectors *X* were built as described, using the n-back CS50→R and CS50→N trials (*n* = 1,2, …, *N* with *N* = 6). In case (2), the regressor vectors *X* were built as described, using the n-back CS100 and CS0 trials (*n* = 1,2, …, *N* with *N* = 6).

To better compare the anticipatory licking from different sessions, the coefficients of each regressor *n* were standardized as follows 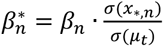 and, for better interpretability, transformed as 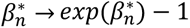. Thus, a positive 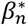 indicates a positive correlation between the anticipatory licks and the *n*-th regressor when the other regressors are constant. Vice-versa, a negative 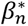 indicates a negative correlation. We performed one regression per session and plotted the average standardized coefficients 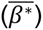 for each animal (n = 88 sessions in 11 animals).

##### Data pre-processing for spike detection

Noise and movement artifacts affecting all recorded channels were reduced by conducting a median subtraction. Therefore, we calculated the median voltage trace of all channels from the same recording site and subtracted the median from each recorded channel. The resulting signal was passed via a band-pass filter (300-5000 Hz, 4th Butterworth filter, built-in MATLAB function). All local maxima crossing a certain amplitude threshold (7.5x of the median absolute deviation of the filtered signal) were identified as spiking events. To prevent a multiphasic spike from being detected multiple times, the minimum distance between threshold crossing peaks was set to 1 ms. If a spiking event was detected on more than one channel of the same tetrode, it was assigned to the timestamp of the highest detected peak. For each cluster, waveforms were obtained by extracting −10 to +21 sampling points around the peak.

##### Data pre-processing for spike sorting

We clustered the detected spiking events using a custom-built graphical user interface in MATLAB developed by A.Koulakov (CSHL). Single-units were separated based on different metrics including peak height or amplitude of the spikes and the respective principal components over channels. When spiking events were predominantly recorded on one channel, the first three principal components of the waveforms were considered. Single-unit quality was quantified using the mlib toolbox by Maik Stüttgen (Version 6, https://de.mathworks.com/matlabcentral/fileexchange/37339-mlib-toolbox-for-analyzing-spike-data). In particular, we estimated cluster quality through refractory period violations (fraction of spikes during the refractory period < 2ms) and waveform variance. Only if clusters had a ratio of refractory period violations to the total number of spikes of less than 2%, they were considered as single-units. In Tu, units with less than 5 Hz baseline firing rate were classified as putative striatal projection neurons and considered for subsequent analysis. Single-units with a firing rate > 5 Hz were excluded as fast spiking neurons from the Tu or from the neighboring anterior ventral pallidum. A total number of 169 putative striatal projection neurons were included in the analysis. In aPC, a total of 486 putative principal units with less than 10 Hz baseline firing rate were included in the analysis.

##### Analysis of single-unit responses

We classified single-units according to their task-related spiking activity. We calculated averaged spike counts for all trial types for the baseline, CS, waiting and US windows (baseline: from −1.5 to −0.5 s; CS: from 0 to 1 s; waiting: from 1 to 2.5 s; US: from 2.7 to 3.7 s relative to odor onset). Single-units were considered as responsive during CS, waiting or US, respectively, if they showed a significant difference compared to baseline (Friedman test, p < 0.05 with Benjamini correction for multiple comparisons on all units tested for each trial type and each task window). Single-units displaying a significant increase in spiking activity were defined as ‘task-excited’ while units displaying a decrease were defined as ‘task-inhibited’ (see **Fig. 4D-F, Fig. 5A-F, Fig. S4D-E and Fig. S5B-C**).

##### Population analysis: the population vector

The session-specific population vector 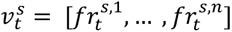 is a vector composed of the firing rate of *n* simultaneously recorded neurons during a time-bin centered at time *t* = 1 … *T*, with *T* total number of time steps per trial. Given the relatively small cell yield per session, session-specific population vectors were concatenated to compose a global population vector 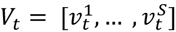. Trials of different sessions were concatenated by matching CS, trial order, and, for CS50, trial outcome. Since, in CS50 trials, reward was delivered with a 0.5 probability, not all sessions had the same number of rewarded CS50 trials. Population vectors for CS50 were, thus, built by selecting the minimum number of trials available among sessions and, in sessions with supernumerary trials, by omitting the last trials.

##### Population analysis: deviation from baseline

To investigate the temporal evolution of aPC and Tu responses to CS and US, we computed the deviation of the population vector from its baseline configuration. At each time step *t*, deviation from baseline was computed as Euclidean distance between the population vector *V*_t_ and the baseline vector *B*. The *B* vector was composed of the firing rate of each unit in the window −2 to −1.25 s before odor onset averaged across all trials. To reduce trial-to-trial variability, such analysis was performed by using the population vectors 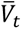, built by grouping population vectors *V*_t_ of consecutive trials in pairs and averaging. This procedure reduces the number of samples available but produces more stable population responses. For visualization, population vectors were computed on bins of 500 ms, moved in time with steps of 125 ms. Statistical tests of changes in population response to different trial types were conducted on distances computed with non-overlapping bins of 250 ms.

##### Population analysis: cross-trial correlation

To assess if the different trial types recruit overlapping sets of units and to quantify the degree of such overlap throughout the trial progression, we computed the Pearson cross-correlation between the population vectors of trials with different trial types. Trials were first grouped according to their CS trial type. Then, for each time step *t*, each vector *V*_t_ was cross-correlated, at turn, with the *V*_t_ vectors of all trials from other CS trial types. For example, to quantify the degree of overlap between the set of units supporting the encoding of CS100 and CS0 during the trial we computed, at every time step *t*, the average Pearson cross-correlation between *V*_t_ of all CS100 trials with *V*_t_ of all CS0 trials. An increase (decrease) in cross-correlation will imply an increase (decrease) in the size of the subpopulation commonly recruited by the two stimuli at a certain time *t*. As above, such analysis was performed by using population vectors 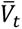, built by grouping population vectors *V*_t_ of consecutive trials into groups of two and averaging. Population vectors were computed on bins of 500 ms, moved in time with steps of 125 ms.

##### Population analysis: population trajectories

To visualize the differences between the temporal evolution of the population vectors in response to different CS and trial types, we averaged the population vectors *V*_t_ across all trials to obtain, for each trial type, a matrix of size N x T. To improve the reconstruction of the neuronal dynamics (Balaguer-Ballester et al., 2011), we applied time embedding on the multivariate time series obtained. We used m = 4 delayed coordinates with a delay constant of 1 bin. Finally, for visualization, we reduced with PCA the space dimensionality to 3.

##### Identification of functional unit-clusters

To identify functional unit-clusters within aPC and Tu populations we used a clustering approach adapted from Cohen et al., 2012. Functional clusters were established according to the similarity in unit responses to different trial types. We first quantified the deviation at time *t* of the response of each unit to its baseline distribution by computing the area under the receiver operating characteristic curve (auROC). The baseline distribution was computed by pooling the firing rate of the unit in the window from −1.8 to −1.4 s before odor onset. The distribution of the unit response at time *t* was computed similarly, but pooling from a window of 500 ms shifted along the trial in steps of 125 ms. auROC values were computed for each unit and each condition - that is CS0, unrewarded CS50, rewarded CS50, CS100 – and concatenated as shown in **Fig. 4G** and **Fig. 5J**. We then performed PCA on the concatenated profiles and applied hierarchical clustering. Hierarchical clustering was performed on the Euclidian distance between vectors comprising of the first 5 PCs of each unit. For aPC and Tu, we used a cutoff in the linkage tree of 0.45 and 0.5, respectively. Such cutoffs were chosen to balance between generality and representativeness of the clusters with respect to their composing units.

##### Complete and distributed coding

To assess if aPC and Tu encoded a monotonic RP in a homogenous or distributed fashion (**Fig. 6, S6**), we tested each unit for monotonic RP coding and single CS dominance. To test monotonic RP coding during CS and the waiting period, we computed the unit firing rate in the CS window (0 to 1 s after odor onset) and waiting period (1 to 2.5 s after odor onset), respectively. Unit responses were computed for each trial *tr* and grouped by CS into *R*0*_tr_*, *R*50*_tr_*, *R*100*_tr_*. Units from the transient-, ramping- and sustained-clusters were labeled as monotonic RP coding if {*R*0*_tr_*} < {*R*50*_tr_*} and *R*50*_tr_*} < {*R*100*_tr_*} when tested with a two-tailed Wilcoxon rank-sum test (p-value < 0.05). If the test for monotonic RP coding failed, units were further tested for CS dominance. CS dominance required that responses to the CS with stronger unit response were significantly higher than those to the CS with the second-strongest response (two-tailed Wilcoxon rank-sum test, p-value < 0.05). For units from the inhibited-clusters, monotonic RP was established by testing {*R*0*_tr_*} > {*R*50*_tr_*} and {*R*50*_tr_*} > {*R*100*_tr_*} while CS dominance required that responses to the CS with strongest task-inhibited response dropped to significantly lower response rates than those to the CS with the second-lowest response.

##### Reward surprise

To test if a unit encoded surprise in receiving an unexpected reward (reward surprise) we tested if the rate jump 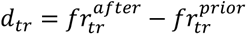 from prior to after US onset was bigger for rewarded CS50 trials than for CS100 trials. To encode surprise we enforced two prerequisites: a) the average rate jump for CS50 rewarded trials had to be positive, 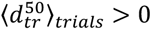 b) the rate jump for CS50 rewarded trials had to be bigger than that for CS100 in a two-tailed Wilcoxon rank-sum test (significance fixed at p-value < 0.05), 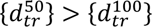. From visual inspection of the activation profile of Tu and aPC units, two characteristic US response profiles emerged, with different latency from US and different duration. Clustering units in transient- or ramping-populations did not segregate the two response profiles. To capture both response types, we computed reward surprise in two distinct trial windows. To capture short-latency short-duration responses, 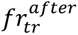 was computed in the window from 2.7 to 3.2 s after odor onset. To capture longer latency responses with longer duration, 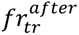 was computed in the window from 3.2 to 4.5 s after odor onset. In both cases, 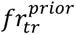 was the firing rate of the unit at trial *tr* in the 500 ms window prior US delivery. All units were tested in both windows and were flagged as encoding reward surprise if significant in either window. The criteria listed here were applied to all units from the transient-, ramping-and sustained-clusters. For units from the inhibited-cluster, reward surprise required: a) 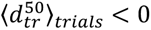; b) 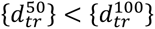.

##### Outcome discrimination

To assess if a unit encoded outcome discrimination, we used a two-tailed Wilcoxon rank-sum test and tested if 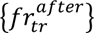 for CS50 rewarded trials was bigger than 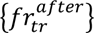 for CS50 unrewarded trials. We did not consider the extremely rare opposite case, thereby focusing on the mechanisms contributing to the computation of the PE. Similar to the test for reward surprise, we tested both short- and long-latency responses and tested units from the task-inhibited cluster using the opposite inequity sign (see ‘Reward surprise’ paragraph).

##### Chance level for monotonic RP coding

We selected all #_discriminative units_ units which showed a different response intensity to the three CS. This was obtained by requiring a p-value < 0.05 when testing the responses of each unit to different pairs of odors with a two-tailed Wilcoxon rank-sum test. Since the number of permutations without repetition of *n* = 3 elements is *n*! = 6, the number of units with a specific order in odor response (e.g. CS100 > CS50 > CS0) expected by chance is #_discriminative units_/6.

### QUANTIFICATION AND STATISTICAL ANALYSIS

#### Functional MRI data

fMRI sessions were analyzed using SPM12 and the SPM12-based Sandwich Estimator toolbox (SwE) at the group-level. Unless otherwise stated, the statistical threshold was set to p < 0.025 (for two-sided testing), false discovery rate (FDR)-corrected for testing of multiple voxels.

#### Behavioral and electrophysiological data

Behavioral and electrophysiological data were analyzed using built-in and custom-made MATLAB routines (Mathworks) and SPSS (IBM). Statistical tests: test statistics, sample size, and multiple comparison corrections were indicated for each performed test and reported either in the relative method sections or in **Table S1**. Graphical visualization: whenever reporting averaged or collective results, the number of units used was reported directly next to the graph (as n); box plots were centered on the median, indicated the 25^th^ and 75^th^ percentiles with solid polygons and extended with whiskers to the most extreme data points not considering outliers, outliers were indicated with circles; average values were always reported with SEM. In all figures, asterisks marked p - values < 0.05 whose exact value was reported in **Table S1**.

## SUPPLEMENTARY FIGURES

**Figure S1.**
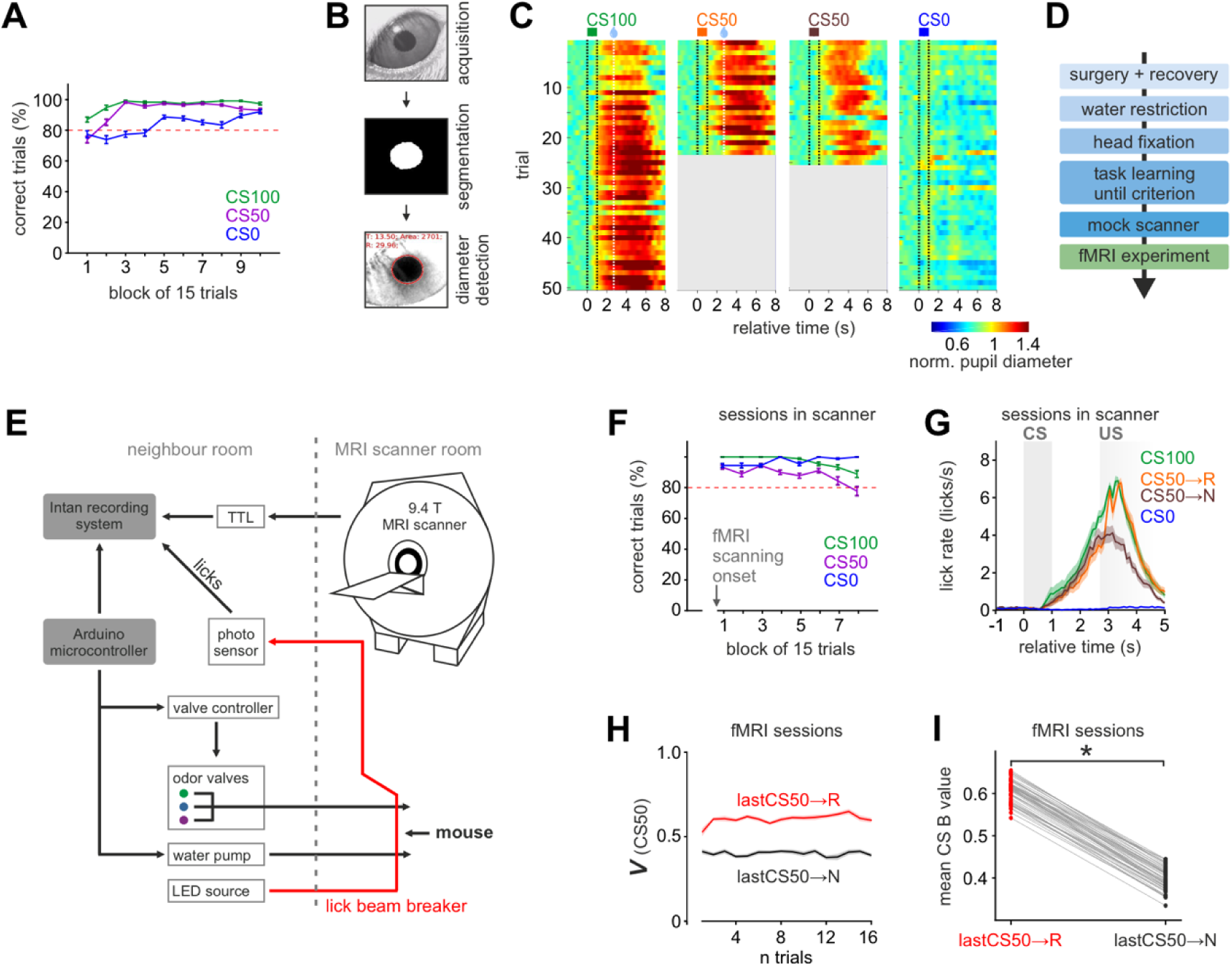
Trained mice display differential anticipatory responses to olfactory stimulus-outcome pairs and dynamic value update by recent outcome-history. Related to Figure 1. (**A**) Percent correct trials for the three CS types in the mouse cohort shown in Fig. 1 (mean ± SEM, n = 69 sessions in 23 animals). One trial block had 5 trials of each type. Mice performed consistently above criterion. (**B**) Pupil image processing. Conversion from raw video to binary frames for pupil segmentation and pupil diameter detection. (**C**) Pupil responses to each trial type from one example session. Color indicates an increase (red) or decrease (blue) from baseline. (**D**) Standard procedure with surgery and training before the fMRI experiments. (**E**) Schematic of the MRI-compatible setup. Odors were delivered through a custom-built olfactometer. Water was delivered by a remote syringe pump. Licks were detected with custom-built optics. The setup was controlled by Arduino microcontrollers. **(F-G)** Performance curves (F) and average lick rates ± SEM (G) for the sessions during fMRI acquisition. (**H**) Same as Fig. 1I but computed with fMRI CS-US sequences. A dynamic update of reward prediction in CS50 trials by the prior cue-specific outcome was also reflected in the TD modeling of the fMRI cohort. Since mice already performed 20-30 trials before fMRI acquisition started and reached a stable performance (F), initial value conditions were set to the reward probabilities associated with each CS. (**I**) Average CS50 value from TD model for lastCS50→R and lastCS50→N (one data point per session) were significantly different depending on the outcome of the previous CS50 trial (Mann-Whitney U test).

**Figure S2.**
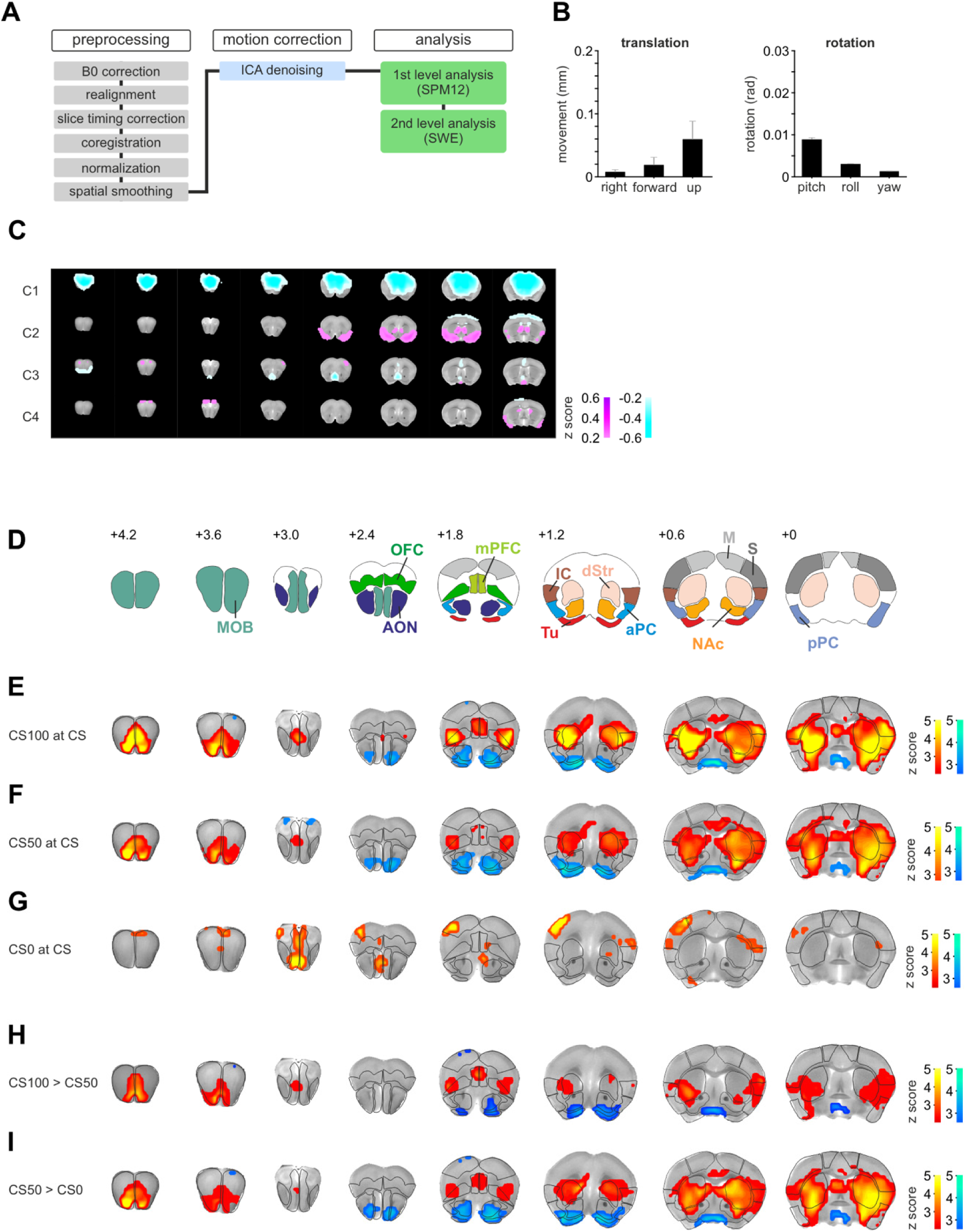
Distributed brain circuits code for monotonic RP and the PE. Related to Figure 2. (**A**) Processing pipeline of fMRI data. See Methods for detailed information. (**B**) Average head motion parameters ± SEM after habituation to the scanner environment for (left) the three translation and (right) the three rotation parameters. Motion parameters of n = 51 sessions in 18 animals were considered. (**C**) Z-statistical maps showing four motion-related independent components. Motion-related components were removed before first level analysis was conducted. (**D**) Same as Figure 2B. ROI definitions based on Paxinos atlas. **(E-G)** Group-level Z-statistical maps for the (E) CS100, (F) CS50, and (G) CS0 regressor (n = 51 sessions in 18 animals). Statistical threshold was set to p < 0.025 false discovery rate (FDR)-corrected, for two-sided testing (as for the other maps unless otherwise indicated). Red colors indicate areas activated by the respective CS events, while blue colors indicate deactivation. Olfactory and striatal areas were primarily recruited. (**H**) Group-level Z-statistical contrast maps for CS100>CS50 used for the RP intersection. The analysis was restricted to the regions associated with *V*(*CS*) from Fig. 2C. (**I**) Same as (H) showing the contrast maps for CS50>CS0.

**Figure S3.**
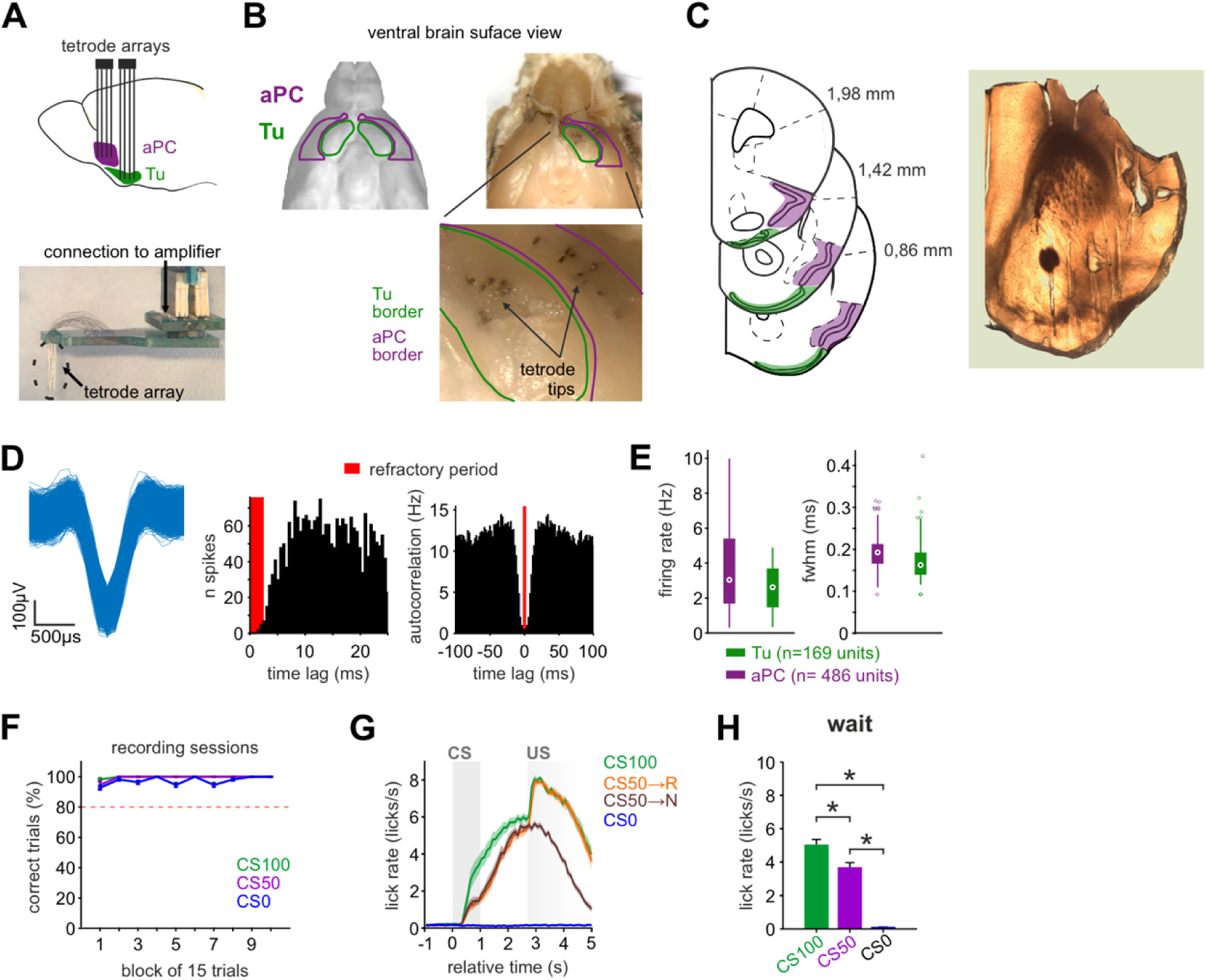
Single-unit recordings in the olfactory tubercle and the anterior piriform cortex. Related to Figure 3. (**A**) Scheme with the anatomical relation between Tu and aPC in sagittal view (top) and an example of a tetrode array connected to the breakout board of the head stage connector (bottom). Tetrode arrays were implanted unilaterally. (**B**) Ventral forebrain view of the mouse brain including ROI definitions. The tetrodes tips are visible in the Tu (green) and aPC (purple) on the ventral surface of the brain. (**C**) Localization of Tu and aPC on different atlas sections (left). Localization from bregma is indicated for each section. Tetrode array tracks are shown in a coronal brain section (right). (**D**) Quality metrics to evaluate spike sorting. Example single unit showing clustered action potential waveforms (left). Only units with less than 2% violations of the refractory spike period (middle) were included. Spike autocorrelation (right). (**E**) Baseline firing rate (left) and full width at half maximum (fwhm) of the action potential (right) of the analyzed single units. (**F**) Performance curves of the mouse cohort used in electrophysiology experiments (mean ± SEM, n = 88 sessions in 11 animals). Only mice were included that performed above criterion. (**G**) Average lick rate ± SEM during different trial types (n = 88 sessions in 11 animals). (**H**) Average lick rate ± SEM in the waiting window differentiated the respective trial types with CS100>CS50>CS0 (one-way ANOVA with Tukey post-hoc comparisons). In the figure: * indicates p < 0.05 (see Table S1 for exact p-values and test details).

**Figure S4.**
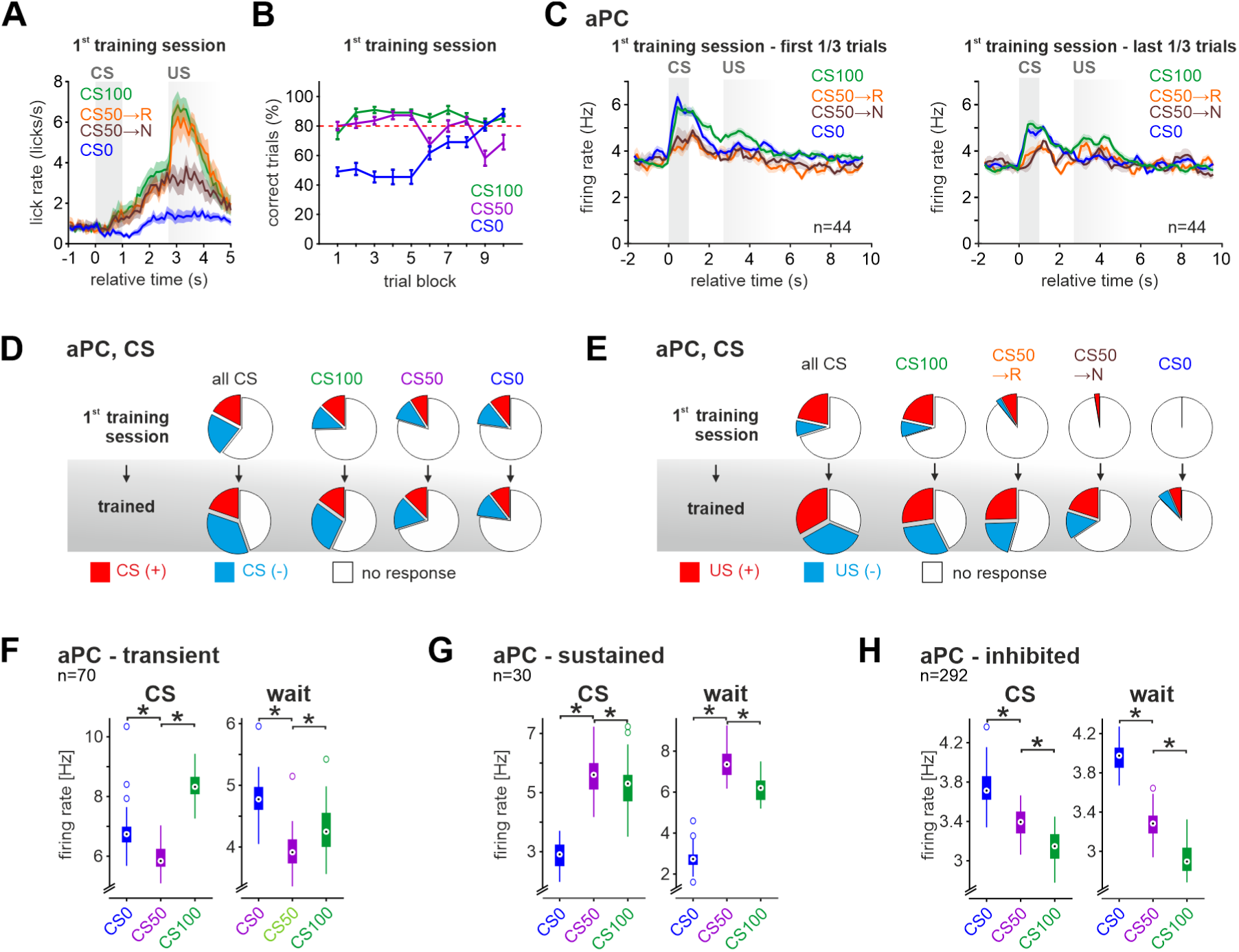
Evolution of task-related responses during training. Related to Figure 4. (**A**) Average licking rate ± SEM split by trial types for the first training sessions (n = 11 sessions in 11 animals). (**B**) Percentages of the correct CS100, CS50 and CS0 trials in first training sessions (mean ± SEM). (**C**) Mean firing rate ± SEM of aPC units recorded during the first training sessions. Displayed mean firing rate during the first 50 trials (left) and the last 50 trials (right) of the session. **(D-E)** The fraction of task-inhibited responses at (D) CS and (E) US increased in aPC with training. **(F-H)** Box plots with median firing rate of the aPC (F) transient-cluster, (G) sustained-cluster and (H) inhibited-cluster during CS and waiting period (one-way ANOVA with Tukey post-hoc comparisons). In the figure: * indicates p < 0.05 (see Table S1 for exact p-values and test details) and n indicates the number of units.

**Figure S5.**
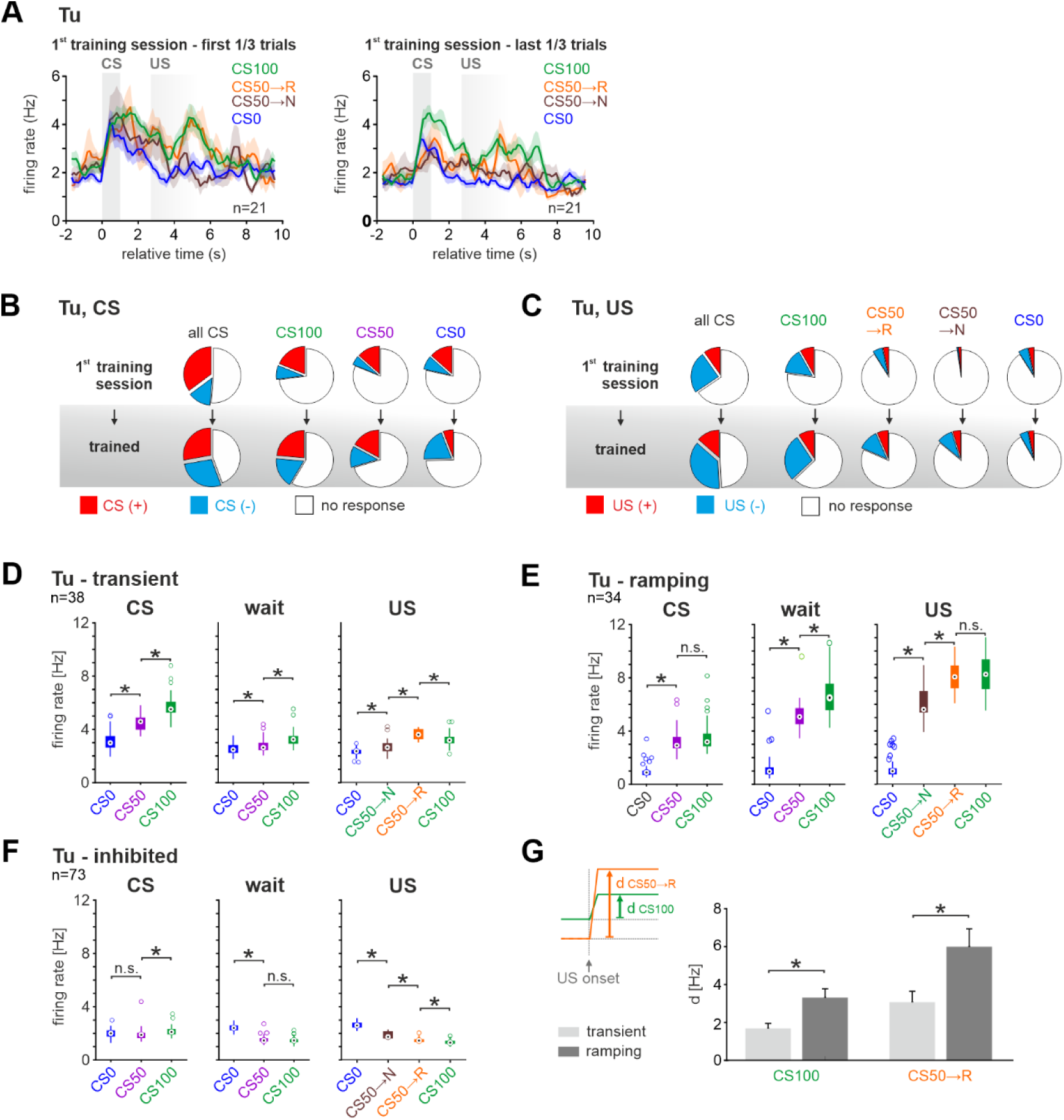
Tu units form transient and ramping task response clusters. Related to Figure 5. **(A)** Same as Fig. S4C for Tu. By the end of the first session, the Tu firing rate started reflecting RP coding by responding more strongly to CS100 than CS50 and CS0. Interestingly, at this stage, ramping activity is not yet seen during the waiting period. **(B-C)** Same as Fig. S4D-E for Tu. Also in Tu, the fraction of task-inhibited responses increased with training for all trial types. **(D-F)** Same as Fig. S4F-H for the Tu (D) transient-cluster, (E) ramping-cluster and (F) inhibited-cluster during CS, waiting period, and US (one-way ANOVA with Tukey post-hoc comparisons). **(G)** Average reward response ± SEM of units from the transient- and ramping-cluster during the CS100 and the rewarded CS50 trials. Reward response was computed as the difference in the firing rate after US from before US. Units from the ramping-cluster had a stronger reward response than those in the transient-cluster regardless of the certainty or not of the reward (CS100 or CS50, respectively) (two-tailed unpaired Wilcoxon rank-sum test). In the figure: * indicates p < 0.05 (see Table S1 for exact p-values and test details) and n indicates the number of units.

**Figure S6.**
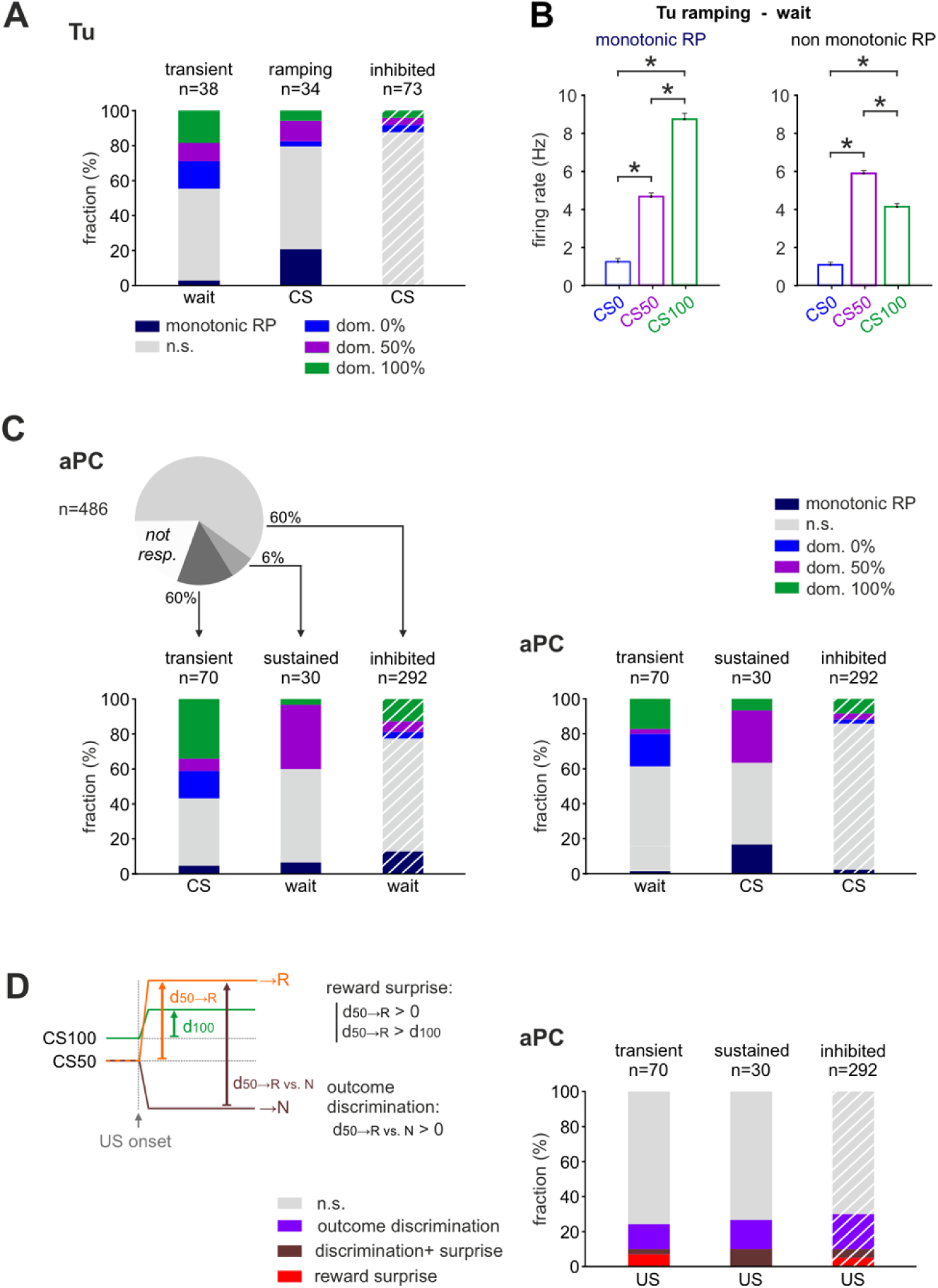
Transient and ramping units differently encode monotonic RP. Related to Figure 6. (**A**) Same as Figure 6A (bottom) but with complementary test windows. Note that coding in the three clusters changed between the CS and the waiting period. (**B**) Same as Figure 6C but for the ramping Tu cluster group. No distributed monotonic RP coding was found in the Tu ramping group during the waiting period. (**C**) Same as Fig. 6A and (A) for aPC. The percentage of units in the three clusters defined in Fig. 4H-J coding individually for monotonic RP or showing a dominant activation for one of the three CS. Right: Complementary test windows. In aPC, only a small fraction of units in the inhibited-cluster encoded monotonic RP during the waiting period (in the task-excited clusters such fraction was below chance). (**D**) Same as Fig. 6D for aPC. Fraction of aPC units in the three clusters defined in Fig. 4H-J encoding reward surprise, outcome discrimination, or both. Note that units from all clusters contributed to some extent to PE coding. In the figure: * indicates p < 0.05 (see Table S1 for exact p-values and test details) and n indicates the number of units.

**Figure S7.**
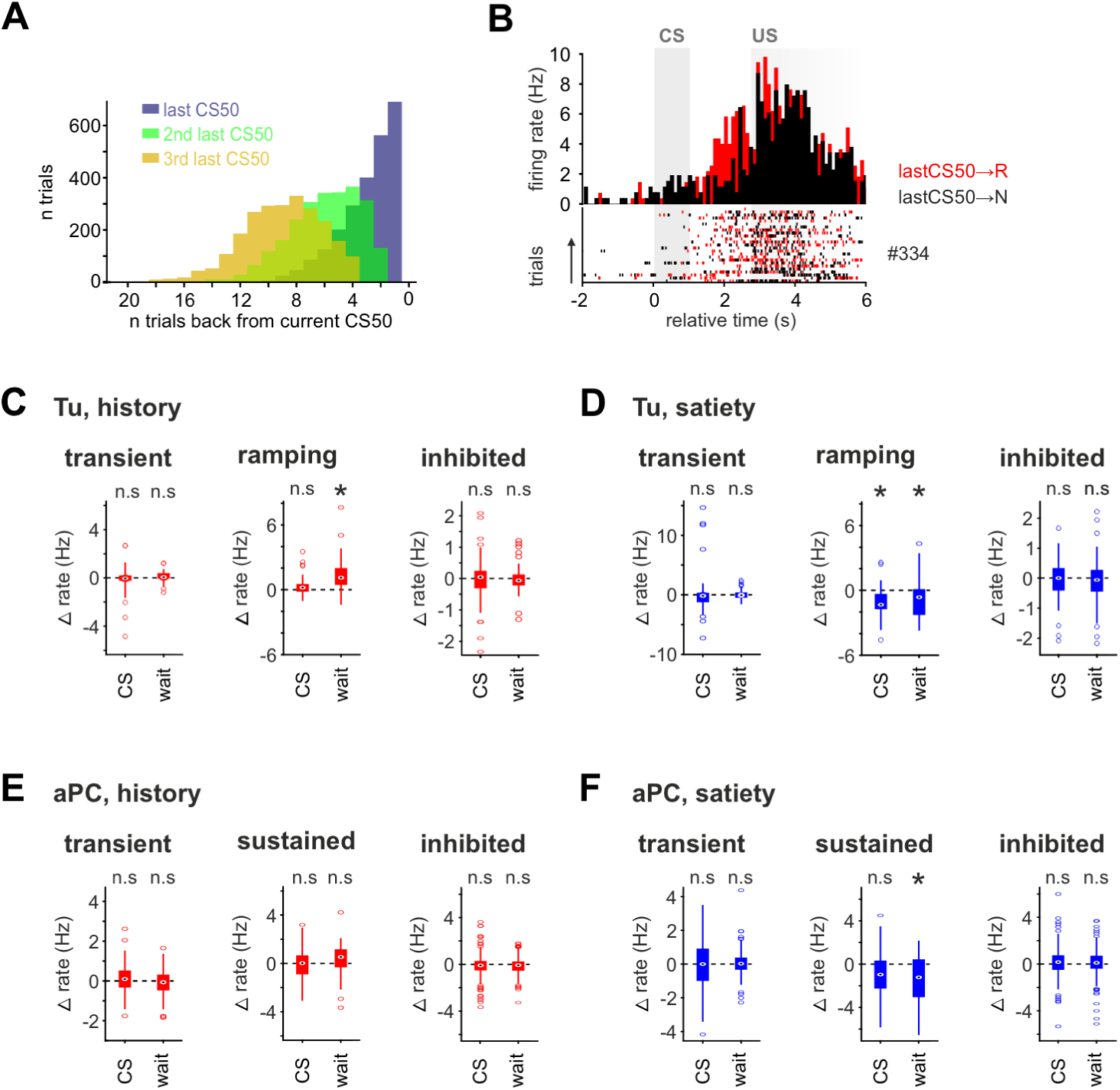
Reward prediction updating through the recent cue-specific outcome-history. Related to Figure 7. (**A**) Histograms showing the distribution of the trial-gap between a CS50 trial and the respective last, second to last, and third to last CS50 trials, in all recording sessions. (**B**) PSTH and raster plot of an example Tu unit from the ramping-cluster coding for outcome-history. **(C-D)** Difference in mean activity for the three major clusters in Tu for outcome-history (C) and satiety (D). One-way repeated measures ANOVA with Greenhouse-Geisser and Bonferroni correction performed for history and satiety separately at CS and during waiting. Note that the ramping-cluster encoded both outcome-history and satiety during waiting. **(E-F)** Same as (C-D) for the aPC clusters. No cluster encoded outcome-history in aPC. In the figure: * indicates p < 0.05 (see Table S1 for exact p-values and test details).

## Notes

### Competing Interest Statement

The authors have declared no competing interest.

